# Functional identification of soluble uric acid as an endogenous inhibitor of CD38

**DOI:** 10.1101/2023.06.03.543541

**Authors:** Shijie Wen, Hiroshi Arakawa, Shigeru Yokoyama, Yoshiyuki Shirasaka, Haruhiro Higashida, Ikumi Tamai

## Abstract

Excessive elevation or reduction of soluble uric acid (sUA) levels has been linked to some of pathological states, raising another subject that sUA at physiological levels may be essential for the maintenance of health. Yet, the fundamental physiological functions and molecular targets of sUA remain largely unknown. Using enzyme assays and *in vitro* and *in vivo* metabolic assays, we demonstrate that sUA directly inhibits the hydrolase and cyclase activities of CD38 via a reversible non-competitive mechanism, thereby limiting nicotinamide adenine dinucleotide (NAD^+^) degradation. CD38 inhibition is restricted to sUA in purine metabolism, and a structural comparison using methyl analogs of sUA such as caffeine metabolites shows that 1,3-dihydroimidazol-2-one is the main functional group. Moreover, sUA at physiological levels prevents crude lipopolysaccharide (cLPS)-induced systemic inflammation and monosodium urate (MSU) crystal-induced peritonitis in mice by interacting with CD38. Together, this study unveils an unexpected physiological role for sUA in controlling NAD^+^ availability and innate immunity through CD38 inhibition, providing a new perspective on sUA homeostasis and purine metabolism.

## Introduction

The evolutionary loss of uricase activity in humans and certain primates completely blocks the degradation of soluble uric acid (sUA), leading to higher physiological levels of sUA than in other mammals (Oda, Satta, Takenaka, & Takahata, 2002; Wu, Muzny, Lee, & Caskey, 1992). Owing to renal reabsorption, sUA is strictly maintained in humans rather than being eliminated as a waste. It has been suggested that sUA functions as an abundant antioxidant (Glantzounis, Tsimoyiannis, Kappas, & Galaris, 2005) and is crucial to maintain blood pressure (Watanabe et al., 2002). Although hyperuricemia may promote the precipitation of monosodium urate (MSU) crystal, an activator of the NLRP3 inflammasome (Martinon, Pétrilli, Mayor, Tardivel, & Tschopp, 2006), resulting in diseases such as gout (Dehlin, Jacobsson, & Roddy, 2020) and kidney stone disease (Howles & Thakker, 2020), numerous studies have indicated the protective potential of sUA in neurodegenerative diseases (Kutzing & Firestein, 2008; Lu et al., 2016; Scott et al., 2002). In addition, abnormal reduction of sUA levels is clinically associated with the risk of various diseases (Crawley, Jungels, Stenmark, & Fini, 2022; Kutzing & Firestein, 2008), including cardiovascular diseases and kidney diseases. Intriguingly, rapid urate reduction in the initiation of therapy even increases gout flares in patients (Becker et al., 2005). Depletion of sUA in uricase-transgenic mice shortens lifespan and promotes sterile inflammation induced by microbial molecules or pro-inflammatory particles (Kono, Chen, Ontiveros, & Rock, 2010; Shi, 2010). Thus, data supports that sUA provides a physiological defense against excessive inflammation, aging, and certain diseases (Álvarez-Lario & Macarrón-Vicente, 2010; Ames, Cathcart, Schwiers, & Hochstein, 1981; Crawley et al., 2022; Cutler et al., 2019; Kutzing & Firestein, 2008; Lai et al., 2017; Linnerz et al., 2022; Ma et al., 2020; Wan et al., 2020; Wen, Arakawa, & Tamai, 2024). However, the underlying molecular basis of sUA physiology remains poorly understood.

To gain insight into target-based physiological functions of sUA, our laboratory utilized magnetic bead-conjugated 8-oxoguanine (8-OG, an sUA analog; technically, sUA cannot be directly conjugated to the beads) and proteomic analysis to screen the potential candidates of sUA-binding proteins. We discovered several binding proteins of 8-OG (unpublished data), including CD38 that was verified by enzyme inhibition (Figure 1A), which raised our interest in exploring the role of CD38 in sUA physiology. CD38 is mainly expressed in immune cells and has type II or type III membrane orientation, with the catalytic domain facing the outside or inside of cells (Hogan, Chini, & Chini, 2019; Lee & Zhao, 2019; Zhao, Lam, & Lee, 2012). It is also found in intracellular membranes or as intracellular and extracellular soluble forms (E. N. Chini, Chini, Espindola Netto, de Oliveira, & van Schooten, 2018; Funaro et al., 1996; Lee & Zhao, 2019; Zielinska, Barata, & Chini, 2004). Functionally, CD38 serves as a hydrolase to degrade nicotinamide adenine dinucleotide (NAD^+^) (Hogan et al., 2019), an essential cofactor for various metabolic reactions that sustain life (Luongo et al., 2020), as well as its precursor nicotinamide mononucleotide (NMN) (C. C. S. Chini et al., 2020), thus regulating inflammation, aging, and various diseases (E. N. Chini et al., 2018; Hogan et al., 2019). Its cyclase catalyzes the synthesis of cyclic ADP-ribose (cADPR) from NAD^+^ to drive calcium mobilization (Lee, 2001; Lee, Walseth, Bratt, Hayes, & Clapper, 1989), which is crucial for social behavior (Jin et al., 2007) and neutrophil recruitment (Partida-Sánchez et al., 2001).

**Figure 1.**
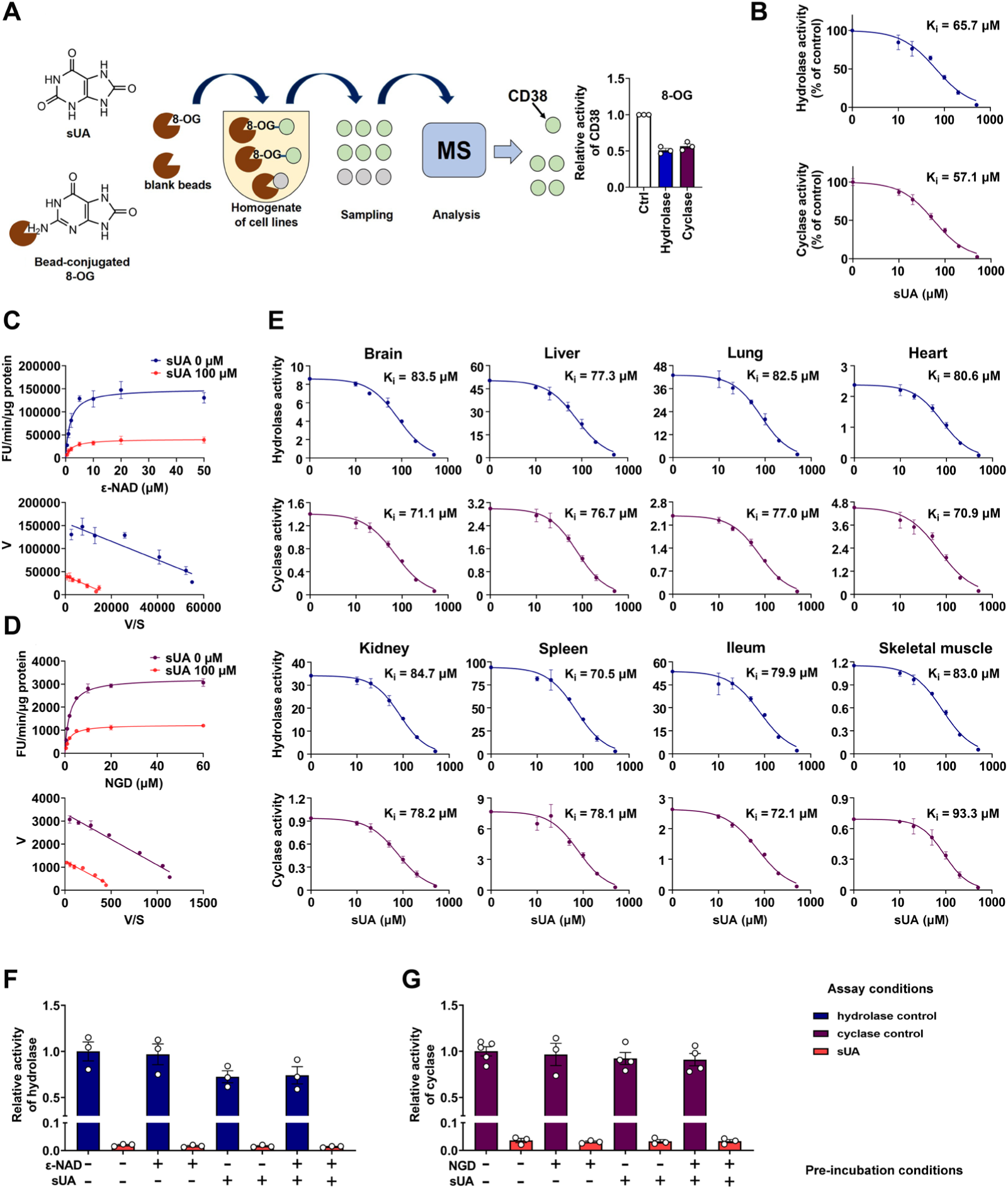
Identification of sUA as an endogenous inhibitor for CD38. **(A)** Preliminary screening of 8-OG binding proteins by mass spectrum (MS)-based proteomics, and effect of 8-OG (50 μM) on CD38 activity (n = 3 experiments/technical replicates). **(B)** Hydrolase and cyclase activities of recombinant human CD38 (hCD38) in the presence of sUA, using nicotinamide 1, N^6^-ethenoadenine dinucleotide (ε-NAD^+^), and nicotinamide guanine dinucleotide (NGD) as substrates, respectively (n = 3 experiments/technical replicates). **(C** and **D)** Effect of different substrate concentrations on sUA inhibition of recombinant hCD38 hydrolase (**C**) and cyclase (**D**) activities (n = 3 experiments/technical replicates). **(E)** Effect of different sUA concentrations on hydrolase and cyclase activities (FU/min/μg protein) in tissues from 8- to 12-week-old WT mice (n = 3 mice). **(F** and **G)** Reversibility of inhibition of recombinant hCD38 hydrolase (**F**) and cyclase (**G**) activities by sUA. After 30-min pre-incubation as indicated, samples were diluted 100-fold in reaction buffer with or without 500 μM sUA for enzyme assay (n = 3-5 experiments/technical replicates). Data are mean ± s.d. (**B**-**E**) or mean ± s.e.m. (**A**, **F**, and **G**).

Here, we demonstrate that sUA at physiological levels directly inhibits CD38 and consequently limits NAD^+^ degradation and excessive inflammation, which defines, for the first time, the physiological functions of sUA via CD38. In addition, we confirm the unique effect of sUA on CD38 in purine metabolism and identify a structural feature for pharmacological inhibition of CD38.

## Results

### sUA is an endogenous, reversible, and non-competitive inhibitor of CD38

To clarify whether CD38 is a direct target for sUA, we investigated the effect of sUA on CD38 activity. We found that sUA directly inhibited the hydrolase and cyclase activities of human (Figure 1B; Figure 1-figure supplement 1A) and murine (Figure 1E) CD38 as a non-competitive inhibitor (Figure 1C and 1D; Figure 1-figure supplement 1B) with a K_i_ in the micromolar range (57.1-93.3 μM), demonstrating its binding to the allosteric sites of CD38. sUA showed comparable inhibitory effects on hydrolase and cyclase, as indicated by similar K_i_. The physiological levels of sUA in humans (about 120-420 μM) (Dalbeth, Gosling, Gaffo, & Abhishek, 2021; Kuwabara et al., 2017) and in several mouse tissues (Figure 1-figure supplement 2) were higher than its K_i_, indicating that human CD38 and murine type III/intracellular CD38 are physiologically inhibited by sUA. The inhibitory effects were reversible (Figure 1F and 1G; Figure 1-figure supplement 1C and 1D), suggesting that sUA dynamically modulates CD38 activity. Interference from endogenous sUA was negligible when using tissues as an enzyme source, because the concentrations in the final reaction buffer were below 1 μM (Figure 1-figure supplement 3).

### CD38 inhibition is restricted to sUA in purine metabolism

Although the structure of sUA is similar to that of other purines, we confirmed its unique effect on CD38 in the major metabolic pathways of purines (Figure 2C). sUA precursors (adenosine, guanosine, inosine, hypoxanthine, and xanthine) and the uricase-catalyzed metabolite (allantoin) hardly inhibited the hydrolase and cyclase activities of human (Figure 2A and 2B; Figure 2-figure supplement 1A) and murine (Figure 2-figure supplement 1B and 1C) CD38. The tested purine concentrations were higher than their physiological levels (Boulieu, Bory, Baltassat, & Gonnet, 1983; Dudzinska, Lubkowska, Dolegowska, Safranow, & Jakubowska, 2010; Eells & Spector, 1983; Traut, 1994). Thus, CD38 inhibition is restricted to sUA, suggesting a specific functional group in sUA.

**Figure 2.**
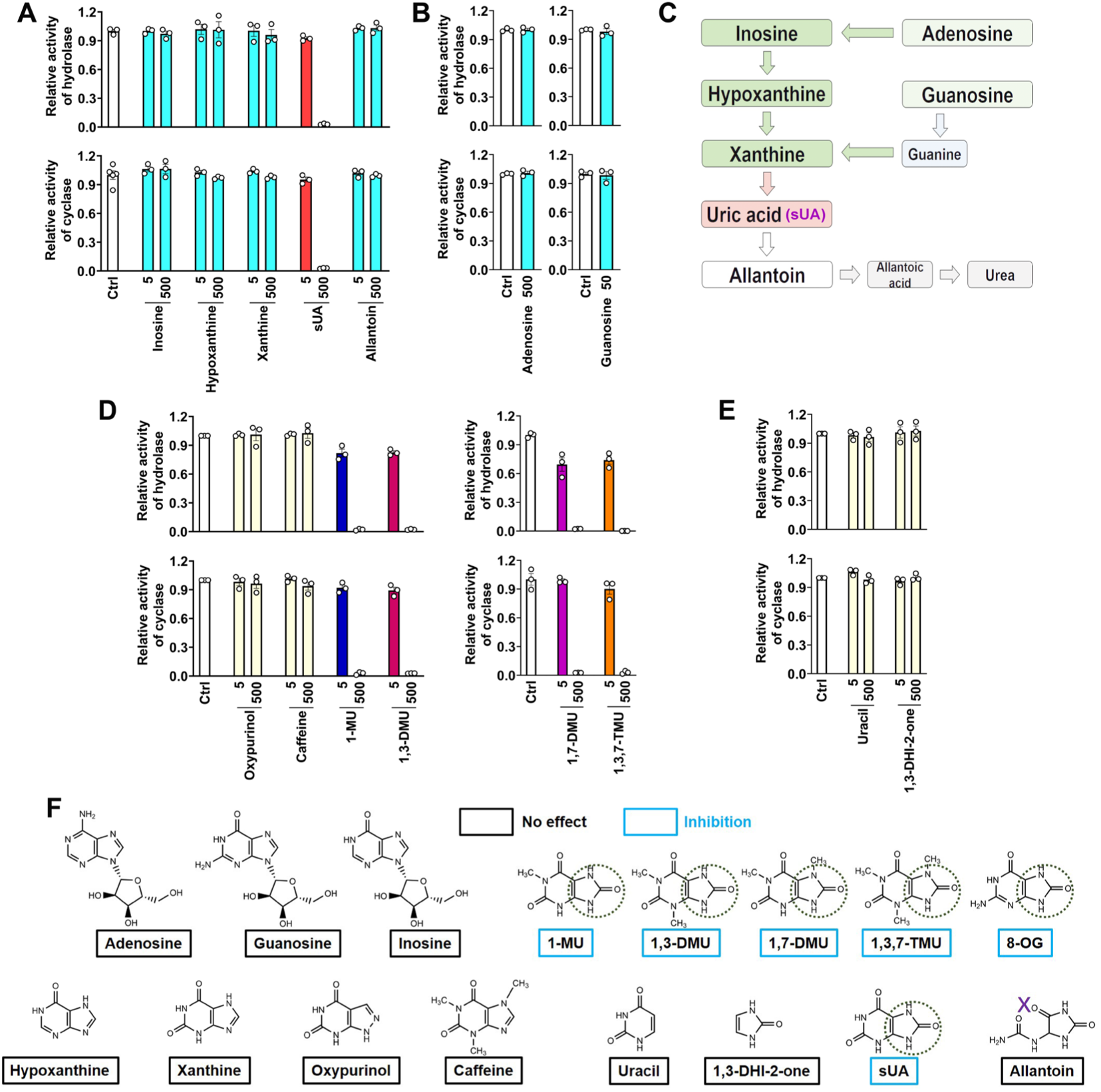
CD38 inhibition is restricted to sUA in purine metabolism. (**A** and **B**) Effect of sUA and its precursors and metabolite on hydrolase and cyclase activities of recombinant hCD38 (n = 3 experiments/technical replicates for each ligand). **(C)** Major pathways of purine metabolism. **(D)** Effect of different analogs on hydrolase and cyclase activities (n = 3 experiments/technical replicates). THP-1 cells were used to detect the effects of oxypurinol, caffeine, 1-MU, and 1,3-DMU on hydrolase activity, recombinant hCD38 was used in the remaining detections. **(E)** Effect of uracil and 1,3-dihydroimidazol-2-one (1,3-DHI-2-one) on hydrolase and cyclase activities of recombinant hCD38 (n = 3 experiments/technical replicates). **(F)** A structural comparison reveals the functional group for CD38 inhibition. The concentrations of all ligands are from 5 to 500 μM. Data are mean ± s.e.m.

To identify the functional group for CD38 inhibition, we tested additional xanthine analogs (oxypurinol and caffeine) and the methyl analogs (caffeine metabolites) of sUA including 1-methyluric acid (1-MU), 1,3-dimethyluric acid (1,3-DMU), 1,7-dimethyluric acid (1,7-DMU), and 1,3,7-trimethyluric acid (1,3,7-TMU). The results showed that only sUA analogs inhibited CD38 activity (Figure 2D). A structural comparison (Figure 2F) indicated that 1,3-dihydroimidazol-2-one (1,3-DHI-2-one) is the main functional group, as all other purines and derivates lacking this group failed to inhibit CD38 activity. In addition, the ring-opening of the uracil group after sUA conversion to allantoin abrogated this inhibitory potential. Uracil or 1,3-DHI-2-one (Figure 2E; Figure 2-figure supplement 1D and 1E) alone did not affect CD38 activity. Therefore, the adjacent uracil-like heterocycles are also essential for CD38 inhibition.

### sUA at physiological levels limits NAD^+^ degradation by directly inhibiting CD38

Next, we explored the effect of sUA on NAD^+^ availability, as CD38 is a key enzyme in degrading NAD^+^ and its precursor NMN (C. C. S. Chini et al., 2020). sUA boosted intracellular NAD^+^ in A549 and THP-1 cells (Figure 3-figure supplement 1) without affecting the activities of nicotinamide phosphoribosyltransferase (NAMPT) and poly (ADP-ribose) polymerase (PARP) (Figure 3-figure supplement 2A and 2C), two key enzymes involved in NAD^+^ synthesis and metabolism, which suggests CD38 as a main target of sUA in regulating NAD^+^ availability. Given that the physiological concentrations of hypoxanthine and xanthine are generally within 20 μM, both NAMPT and PARP might be not affected by purine metabolism under physiological conditions (Figure 3-figure supplement 2B and 2D). Thus, we further investigated whether CD38 mediates the effect of sUA on NAD^+^ degradation. Short-term and moderate “sUA-supplementation” model was constructed by gavage of inosine and oxonic acid (OA, a uricase inhibitor with a K_i_ in the nanomolar range (Fridovich, 1965)) in mice with “natural hypouricemia” (Figure 3D; Figure 3-figure supplement 5C). The plasma sUA levels (around 120 μM) in our models were close to the minimum physiological concentrations in humans but were markedly lower than that in other long-term and hyperuricemia-associated disease models in rodents, which enabled us to evaluate the physiologically inhibitory effect of sUA on CD38 activity. Although OA was a weak inhibitor of CD38 with a IC_50_ in the millimolar range (Figure 3-figure supplement 3A and 3B), it seemed unlikely to affect CD38 activity in our models because of the incomplete inhibition of uricase, as evidenced by plasma sUA (Figure 3-figure supplement 3C). Although short-term administration of OA alone failed to increase plasma sUA levels, we used it as the background for metabolic studies in mice to exclude potential interference.

**Figure 3.**
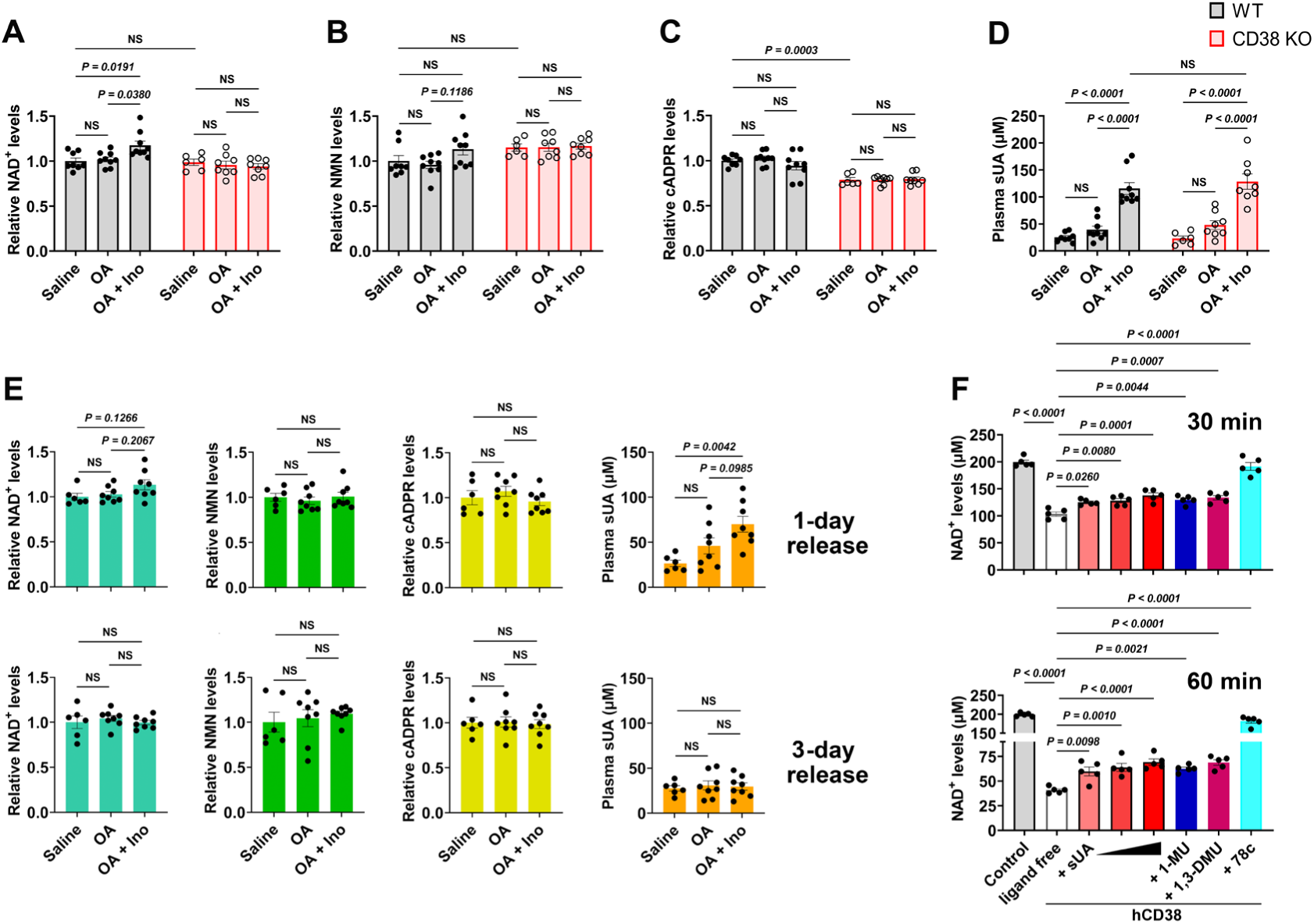
sUA physiologically limits NAD^+^ degradation via CD38 inhibition. WT and CD38 KO mice (10- to 12-week-old) received oral administration of saline, OA, or OA plus inosine (Ino) twice (1-day moderate sUA supplementation). **(A-D)** Effect of 1-day sUA supplementation on whole blood NAD^+^ (**A**), NMN (**B**), cADPR (**C**), and plasma sUA (**D**) levels in WT and CD38 KO mice (WT-Saline: n = 8 mice, WT-OA: n = 9 mice, WT-OA + Ino: n = 9 mice, KO-Saline: n = 6 mice, KO-OA: n = 8 mice, KO-OA + Ino: n = 8 mice). **(E)** Effect of 1-day or 3-day release on whole blood NAD^+^, NMN, cADPR, and plasma sUA levels in WT mice that received 1-day supplementation (WT-Saline: n = 6 mice, WT-OA: n = 8 mice, WT-OA + Ino: n = 8 mice). **(F)** Effect of sUA (100, 200, or 500 μM) and other ligands (analogs at 500 μM, 78c, a CD38 inhibitor, at 0.5 μM) on NAD^+^ degradation by recombinant hCD38 (n = 5 independent samples). Data are mean ± s.e.m. Significance was tested using 2-way ANOVA (**A-D**), Kruskal-Wallis test or 1-way ANOVA with Tukey’s multiple comparisons test (**E** and **F**). NS, not significant. Statistic difference (**A**-**D**) between OA and OA + Ino groups in WT or KO mice (saline alone group excluded) was also analyzed by unpaired two-sided t-test or Mann Whitney test; WT mice: *P* = 0.0056 (**A**), 0.0351 (**B**), or *P* < 0.0001 (**D**); KO mice: *P* = 0.0003 (**D**).

One-day or 3-day moderate sUA supplementation slightly but significantly increased whole blood NAD^+^ levels in wild-type (WT) mice but not in CD38 knockout (KO) mice (Figure 3A and 3D; Figure 3-figure supplement 5C and 5D). Similar results were also observed under inflammatory conditions (Figure 3-figure supplement 4A and 4B; Figure 4-figure supplement 4D and 4E), and OA or inosine alone had no interference (Figure 3-figure supplement 4E-4H). This indicates that CD38 mediates, at least in part, the suppressive effect of sUA on NAD^+^ degradation. The suppression of NAD^+^ degradation by the sUA-CD38 interaction was validated by a decrease in cADPR production under inflammatory conditions (Figure 3-figure supplement 4D) but not under non-inflammatory conditions (Figure 3C; Figure 3-figure supplement 5F), possibly because of physiological compensation via other cADPR synthases. However, sUA had a minor effect on NMN levels *in vivo* (Figure 3B; Figure 3-figure supplement 4C; Figure 3-figure supplement 5E; Figure 4-figure supplement 4F), likely due to the rapid conversion of NMN to NAD^+^ (Mills et al., 2016). Notably, WT and CD38 KO mice showed similar NAD^+^ baselines in whole blood, suggesting that the metabolic background has been reprogrammed in CD38 KO mice; for instance, sirtuins with higher activity in CD38 KO mice (Camacho-Pereira et al., 2016) may consume more NAD^+^. Moreover, sUA release *in vivo* showed that both plasma sUA and whole blood NAD^+^ gradually returned to their initial levels (Figure 3E), which confirmed the reversible regulation of NAD^+^ availability by sUA and excluded potential interference from epigenetic regulation of other bioconversion pathways of NAD^+^. Metabolic assays using recombinant hCD38 further verified that increased NAD^+^ availability is mediated by direct CD38-sUA interaction (Figure 3F).

**Figure 4.**
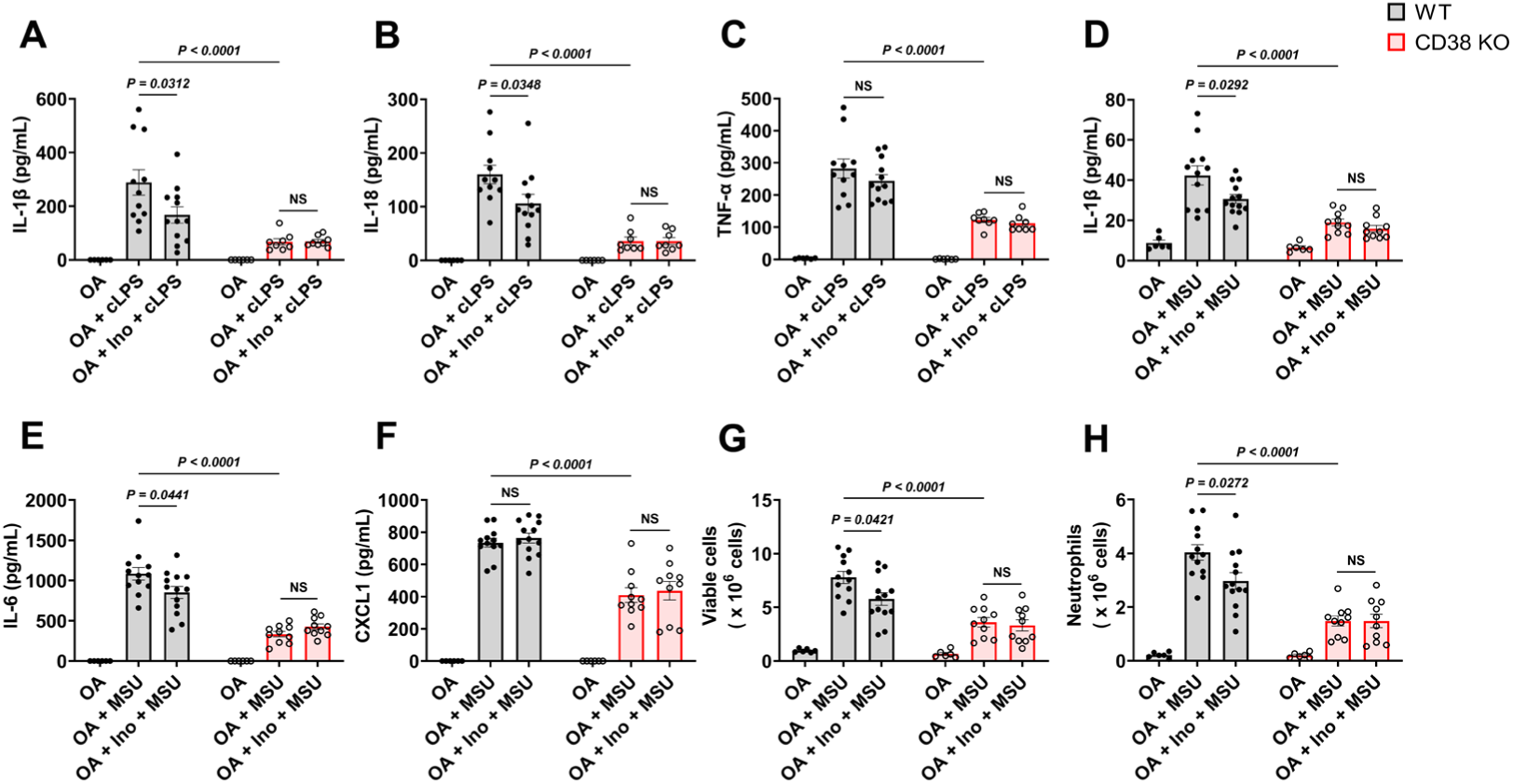
sUA physiologically prevents excessive inflammation by interacting with CD38. WT and CD38 KO mice (10- to 12-week-old) received 1-day moderate sUA supplementation, plasma sUA was increased to the minimum physiological levels of humans in OA plus inosine (Ino) groups. Then, the mice were stimulated with cLPS (2 mg/kg) or MSU crystals (2 mg/mouse) for 6 h. **(A-C)** Effect of sUA at physiological levels on serum levels of IL-1β (**A**), IL-18 (**B**), and TNF-α (**C**) in mice with cLPS-induced systemic inflammation (WT-OA: n = 6 mice, WT-OA + cLPS: n = 11 mice, WT-OA + Ino + cLPS: n = 12 mice, KO-OA: n = 6 mice, KO-OA + cLPS: n = 8 mice, KO-OA + Ino + cLPS: n = 8 mice). **(D-H)** Effect of sUA at physiological levels on IL-1β (**D**), IL-6 (**E**), and CXCL1 (**F**) levels and recruitment of viable cells (red blood cells excluded) (**G**) and neutrophils (**H**) in peritoneal lavage fluid from the mice with MSU crystal-induced peritonitis (WT-OA: n = 6 mice, WT-OA+MSU: n = 12 mice, WT-OA + Ino + MSU: n = 13 mice, KO-OA: n = 6 mice, KO-OA+MSU: n = 10 mice, KO-OA + Ino + MSU: n = 10 mice). Data are mean ± s.e.m. Significance was tested using 2-way ANOVA with Tukey’s multiple comparisons test. NS, not significant. Statistic difference between OA + cLPS/MSU and OA + Ino + cLPS/MSU groups in WT or KO mice (OA alone group excluded) was also analyzed by unpaired two-sided t-test (with Welch’s correction when applicable) or Mann Whitney test; WT mice: *P* = 0.0398 (**A**), 0.0383 (**B**), 0.0407 (**D**), 0.0417 (**E**), 0.0222 (**G**), or 0.0205 (**H**).

In contrast, the tissue levels of NAD^+^ and NMN were not increased in 1-day and 3-day sUA-supplementation models, probably because the tissue uptake of sUA was physiologically saturated before treatment (Figure 3-figure supplement 5A and 5B). The NAD^+^ levels in the brain and heart appeared to be elevated in 7-day model (Figure 3-figure supplement 5G-5I). Thus, the sUA-CD38 interaction in the circulation may indirectly increase tissue NAD^+^ availability. Our models within 7 days did not show pro-inflammatory potential (Figure 3-figure supplement 6), but we did not prolong the treatment time because long-term administration of OA might induce renal damage with concomitant inflammation related to cell death.

Considering the rapid conversion of NMN to NAD^+^ *in vivo*, we assessed the effect of sUA on extracellular NMN degradation *in vitro*. sUA directly inhibited NMN degradation by recombinant hCD38 (Figure 3-figure supplement 7A). Whereas sUA or another CD38 inhibitor failed to boost intracellular NAD^+^ in primed WT bone marrow-derived macrophages (BMDMs) treated with NMN (Figure 3-figure supplement 7B). This is likely because another NAD^+^-consuming enzyme, PARP1, is activated (Minhas et al., 2019) and the NMN transporter expression is very low in BMDMs (C. C. S. Chini et al., 2020). Importantly, sUA increased extracellular NMN availability in WT BMDMs but not in CD38 KO BMDMs (Figure 3-figure supplement 7C and 7D). The addition of recombinant hCD38 restored the suppressive effect of sUA on extracellular NMN degradation in CD38 KO BMDMs (Figure 3-figure supplement 7E). Accordingly, sUA may also increase NAD^+^ availability by inhibiting CD38-medaited NMN degradation.

### sUA physiologically prevents excessive inflammation by interacting with CD38

NAD^+^ is crucial for the activity of sirtuins that limit the NLRP3 inflammasome activation (He et al., 2020; Misawa et al., 2013), and cADPR may regulate calcium signaling to promote cytokine production (Murakami et al., 2012; Zeidler, Kashyap, Hogan, & Chini, 2022). Therefore, CD38 plays a key role in inflammation via NAD^+^ metabolism (Piedra-Quintero, Wilson, Nava, & Guerau-de-Arellano, 2020). We confirmed the role of CD38 in the NLRP3 inflammasome activation. CD38 KO reduced IL-1β release driven by several inflammasome activators and the fungal component zymosan in primed BMDMs (Figure 4-figure supplement 1A). However, pre-incubation with sUA at physiological levels hardly suppressed inflammasome activation in primed WT BMDMs (Figure 4-figure supplement 1B). sUA uptake was very low in primed WT BMDMs (Figure 4-figure supplement 1D), suggesting a crucial role of intracellular sUA in regulating inflammasome activation *in vitro*. However, extracellular sUA may inhibit CD38-mediated NMN degradation to increase NAD^+^ levels *in vivo*, thus limiting excessive inflammation via sirtuins signaling (Misawa et al., 2013). In fact, we noticed that sUA pre-incubation moderately limited IL-1β release in primed THP-1 cells (Figure 4-figure supplement 2A), and CD38 blockade abrogated this effect without reducing sUA uptake (Figure 4-figure supplement 2B and 2C), which suggests CD38 as a key mediator in sUA immunosuppression. Notably, sUA did not induce IL-1β release in primed WT BMDMs and THP-1 cells (Figure 4-figure supplement 1C; Figure 4-figure supplement 2A). These results argue against the pro-inflammatory potential of sUA (Joosten, Crişan, Bjornstad, & Johnson, 2020) that has been challenged by the improper preparation of aqueous solution (MSU crystal precipitation) in basic studies (Alberts et al., 2019; Ma et al., 2020). We also confirmed that long-term storage at 4 ℃ promoted crystal precipitation in high-concentration sUA stock solutions (Figure 4-figure supplement 3).

To reveal the role of sUA-CD38 interaction in regulating inflammation and innate immunity, at first, we stimulated WT and CD38 KO mice with crude lipopolysaccharide (cLPS) after 1-day sUA supplementation. OA was used as the background in mice to exclude its interference. Plasma sUA at the minimum physiological levels of humans (OA plus inosine group) suppressed cLPS-induced production of serum IL-1β and IL-18 in WT mice without affecting inflammasome-independent TNF-α levels, and the suppressive effects were abrogated in CD38 KO mice (Figure 4A-4C), demonstrating that sUA physiologically limits cLPS-induced systemic inflammation via CD38. sUA immunosuppression may be partially mediated by increased NAD^+^ levels and decreased cADPR production in whole blood after sUA-CD38 interaction (Figure 3-figure supplement 4B and 4D). sUA hardly limited high-dose cLPS-induced systemic inflammation (Figure 4-figure supplement 4A-4C), although plasma sUA and whole blood NAD^+^ levels were increased (Figure 4-figure supplement 4D and 4E). In addition, high-dose cLPS, with or without sUA supplementation, did not affect cADPR levels (Figure 4-figure supplement 4G), suggesting some CD38-independent inflammatory mechanisms.

Recently, crystal-free hyperuricemia has been shown to rapidly inhibit neutrophil migration (Ma et al., 2022), suggesting that some biological targets directly mediate sUA immunosuppression (Lowell, 2022). CD38 is crucial for neutrophil recruitment (Partida-Sánchez et al., 2001), we therefore investigated the effect of sUA on MSU crystal-induced peritonitis via CD38. After 1-day sUA supplementation, plasma sUA at the minimum physiological levels of humans (OA plus inosine group) inhibited the recruitment of viable cells and neutrophils and reduced the production of IL-1β and IL-6 in WT mice but not in CD38 KO mice (Figure 4D, 4E, 4G, and 4H), which verified the involvement of CD38 in the suppressive effect of sUA on inflammation and innate immunity. CXCL1 production (Figure 4F) was not affected by sUA, indicating that the sUA-CD38 interaction may inhibit circulating immune cells in response to chemokines. Moreover, sUA may interfere with the interaction between CD38 and adhesion molecules such as CD31(Deaglio et al., 2010; Deaglio et al., 1998) to suppress immune cell migration. Intriguingly, intact IL-8 signaling has been reported in CD38 KO mice (Partida-Sánchez et al., 2001), and sUA at a high concentration (595 μM) inhibits neutrophils in response to IL-8 (Ma et al., 2022). Whereas we did not observe decreased neutrophil recruitment by sUA in CD38 KO mice, likely because plasma sUA levels (around 120 μM) in our models are insufficient to affect IL-8 signaling. In addition, we cannot exclude the possibility that sUA at supraphysiological levels interacts with other targets to regulate IL-8 signaling and β2-integrin, which requires further investigation in the future.

Inosine, an sUA precursor, may also exhibit anti-inflammatory effect by interacting with adenosine receptors (Gomez & Sitkovsky, 2003). Scott et al. reported that after oral administration, serum inosine concentrations slightly increased and then rapidly returned to the initial levels in mice within 2 h (Scott et al., 2002). We stimulated the mice with cLPS or MSU crystals 2 h after the second treatment, suggesting the negligible contribution of inosine. To verify this and exclude the potential contribution of OA, we evaluated the effects of 1-day treatment of inosine or OA on inflammation *in vivo*. The results showed that OA or inosine alone hardly affected cLPS-induced systemic inflammation (Figure 4-figure supplement 5A-5C) and MSU crystal-induced peritonitis (Figure 4-figure supplement 5D-5H), as well as plasma sUA levels and whole blood NAD^+^ metabolism under inflammatory conditions (Figure 3-figure supplement 4E-4H). Thus, these results suggested that sUA at physiological levels limits innate immunity to avoid excessive inflammation by interacting with CD38.

## Discussion

In the present study, we unveiled CD38 as a direct physiological target for sUA and thus defined its fundamental physiological functions in the regulation of NAD^+^ availability and innate immunity, which promotes understanding of the molecular basis of sUA physiology as well as providing an important clue to explore the potential impact of abnormal sUA levels (independent of MSU crystals) on health and disease.

It has been reported that NAD^+^ decline contributes to inflammation, aging-related dysfunction, and multiple diseases, including hearing loss, obesity, diabetes, kidney diseases, and cardiovascular diseases, in murine models (Hogan et al., 2019) and possibly even in humans (Rajman, Chwalek, & Sinclair, 2018). Accordingly, NAD^+^ boosting by CD38 inhibitors has been a promising therapeutic strategy (C. C. S. Chini et al., 2020; E. N. Chini et al., 2018; Escande et al., 2013; Hogan et al., 2019; Peclat, Shi, Varga, & Chini, 2020; Tarragó et al., 2018). We discovered a structural feature for pharmacological inhibition of CD38 based on sUA analogs such as caffeine metabolites. Indeed, sUA and its analogs have similar functions, such as antioxidant property (Nishida, 1991) and neuroprotective effects (Haberman et al., 2007), supporting that they share a functional group, 1,3-DHI-2-one, that interacts with the same targets, including but not limited to CD38. Importantly, our results support that sUA at physiological levels limits CD38 activity to maintain NAD^+^ availability, providing the molecular basis for sUA preventing NAD^+^ decline-associated senescence and diseases. sUA levels seem to increase with age (Iwama, Kondo, Shimokado, Maruyama, & Ishigami, 2012; Kuzuya, Ando, Iguchi, & Shimokata, 2002), raising a possibility that sUA elevation within physiological range is a compensatory response to aging in organisms. Given that CD38 expression is relatively low in young mice (Camacho-Pereira et al., 2016), aged mice would be more appropriate to test this hypothesis. The apparent K_i_ values imply that CD38 is completely inhibited by sUA in blood or in some tissues under physiological conditions; however, this might not be the case because other endogenous substance may also regulate CD38 activity (E. N. Chini, de Toledo, Thompson, & Dousa, 1997; Dogan et al., 2002; Hara-Yokoyama et al., 1996). In spite of this, altered sUA levels are able to indicate the changes in the activities of CD38 and other molecular targets of sUA, which partially explains the correlation between sUA homeostasis disruption and disease risk. For instance, abnormal reduction of sUA levels such as hypouricemia may result in higher CD38 activity in the circulation due to the reversible inhibitory effect of sUA, thus negatively influencing health as well as increasing the risk of certain diseases. However, it does not mean that higher sUA levels are better, because excessive elevation of sUA levels promotes the precipitation of MSU crystals. In addition, we cannot exclude the possibility that long-term and crystal-free hyperuricemia (> 420 μM) in humans may overly modulate additional unknown targets, especially in CD38-negative cells, thereby partially covering the CD38-mediated physiological functions of sUA. To identify more molecular targets for sUA, drug affinity responsive target stability (DARTS) (Lomenick et al., 2009) and cellular thermal shift assay (CETSA) (Martinez Molina et al., 2013), which detect the direct interactions between binding proteins and their ligands, may serve as alternative strategies.

Another unexpected observation is the restrictive effect of sUA at physiological levels on excessive inflammation, which is complementary to several recent studies regarding the immunosuppressive effect of crystal-free hyperuricemia in the host (Ma et al., 2020; Ma et al., 2022). Similar to itaconate, an inhibitor of the NLRP3 inflammasome (Hooftman et al., 2020), sUA production increases in activated macrophages (Ives et al., 2015). It has been reported that intracellular sUA reduction promotes certain inflammatory responses (Ives et al., 2015; Kono et al., 2010), hence sUA at physiological levels may function as an endogenous regulator of inflammasomes to avoid excessive inflammation by inhibiting CD38 activity. On the other hand, chemical phase transition (crystal precipitation) is essential for sUA as a danger signal to trigger immune responses (Alberts et al., 2019; Ives et al., 2015; Ma et al., 2020; Ma et al., 2022; Shi, 2010; Shi, Evans, & Rock, 2003). We and another laboratory demonstrated that MSU crystals upregulate CD38 to promote inflammatory responses in primed macrophages (Wen, Arakawa, & Tamai, 2021; Yan, 2021). A recent study confirmed CD38 activation in gout patients (larger-scale analyses are still required) and further validated the role of CD38 in experimental models of gouty inflammation induced by MSU crystals (Alabarse et al., 2024). Importantly, in this study, we showed that CD38 mediates the opposite effects of sUA (even at physiological levels) and MSU crystals on inflammation and innate immunity. Therefore, a sudden and rapid reduction of the circulating sUA levels by high-dose urate-lowering medications may disrupt the immune balance by rapidly releasing CD38 activity before MSU crystal dissolution (Figure 4-figure supplement 6), resulting in the increased risk of gout attack in patients (Becker et al., 2005; Wen et al., 2024), a well-known paradox in gout therapy. Several clinical studies even observed that blood sUA levels decrease during gout flares, and the resolution of gouty inflammation is coming with a recovery of sUA levels (Logan, Morrison, & McGill, 1997; Urano et al., 2002). Given the distinct effects of sUA and MSU crystals on certain targets, intracellular and/or extracellular microcrystals should be excluded when evaluating the pathological role of sUA at high levels in relevant studies. In addition, our findings also provide biological evidence for the neuroprotection of sUA. It has been reported that CD38 is involved in neurodegenerative diseases (Blacher et al., 2015; Meyer et al., 2022); thus, sUA in central nervous system (CNS) tissues may directly inhibit CD38 activity to limit neuroinflammation and the progression of such diseases.

Previously, a physiological medium containing sUA was shown to inhibit UMP synthase to reshape cellular metabolism *in vitro* (Cantor et al., 2017). We identified a unique sUA-CD38 interaction in this study, highlighting the physiologically essential role of sUA as a purine metabolite in sustaining life. Accumulated data suggests a remarkable overlap between the effects of sUA (Álvarez-Lario & Macarrón-Vicente, 2010; Ames et al., 1981; Cutler et al., 2019; Kutzing & Firestein, 2008; Lai et al., 2017; Linnerz et al., 2022; Lu et al., 2016; Ma et al., 2020; Ma et al., 2022; Scott et al., 2002; Wan et al., 2020) and CD38 inhibition/KO (Blacher et al., 2015; E. N. Chini et al., 2018; Hogan et al., 2019; Meyer et al., 2022) in counteracting inflammation, aging, and certain diseases, which also strongly supports our current findings. It should be noticed that we used an exogenous compound as the background in mice to mimic the deficiency of uricase in humans, and tissue sUA levels were unchanged after sUA supplementation, suggesting some of the limitations in our models when evaluating the physiological relevance. Therefore, it is important to extend these studies to global sUA-depletion models in primates or uricase-transgenic mice. Moreover, the commercial recombinant hCD38 without a transmembrane region was partially used in this study, we cannot exclude the potential interaction between sUA and the transmembrane region of CD38, although comparable K_i_ values were obtained in crude enzymes containing full-length CD38. In addition to identification of the allosteric sites of CD38, the crystal structure of the active full-length CD38-sUA complex should be also captured using cryo-electron microscopy to elucidate the inhibitory mechanisms in the future. However, the present study clearly demonstrated that sUA at physiological levels directly inhibits CD38 and consequently limits NAD^+^ degradation and excessive inflammation, suggesting that sUA is crucial for the physiological defense in humans against aging and diseases.

## Materials and methods

### Reagents

Reagents and other materials used in this study are described in Key Resources Table.

### Animals

Male and female ICR (Institute of Cancer Research of the Charles River Laboratories, Inc., Wilmington, MA, USA) mice were initially purchased from Japan SLC, Inc. CD38 KO mice (ICR strain) were generated using the CRISPR/Cas9 method as previously described (Ichinose et al., 2019). WT and CD38 KO mice were kept and bred at the Experimental Animal Center of Kanazawa University (Takara-machi campus). For animal experiments, all the mice (sex and age as indicated in respective figure legends) were transferred a week in advance and housed in the animal room of Research Center of Child Mental Development under standard conditions (24 °C; 12 h light/dark cycle, lights on at 8:30 am) with standard chow and water provided *ad libitum*. Male and female mice were separated after weaning and were equally used for experiments except when specified. In each experimental group, the mice were from different biological mothers. All animal experiments were approved by the Institutional Animal Care and Use Committee at Kanazawa University (AP-214243), and were performed in accordance with ARRIVE and the local guidelines.

### Cell culture

THP-1 and A549 cells were cultured in RPMI-1640 containing 10% FBS and 1% penicillin/streptomycin. Bone marrow cells and bone marrow-derived macrophages were maintained in RPMI-1640 containing 10% FBS, 1% penicillin/streptomycin, and 50 μM 2-mercaptoethanol in the presence or absence of macrophage colony stimulating factor (M-CSF).

### Measurement of CD38 activity

The hydrolase activity of CD38 was measured according to a previous report (de Oliveira, Kanamori, Auxiliadora-Martins, Chini, & Chini, 2018) with minor modifications. CD38 hydrolase activity was measured using 50 μM ε-NAD^+^ as a substrate in hydrolase reaction buffer (250 mM sucrose, 40 mM Tris-HCl, pH 7.4). Briefly, cells or tissues were directly homogenized in blank reaction buffer on ice, and recombinant hCD38 was diluted in blank reaction buffer for subsequent assays. The tissue homogenates were centrifuged to collect the supernatant for enzyme assays. The loading volume of enzyme was 4 to 30 μL, the total volume of the reaction system was 3 mL. To detect enzyme inhibition, except when specified, the ligands were directly dissolved in hydrolase reaction buffer before pH adjustment, after which, the pH was immediately adjusted to 7.4. All reaction buffers, with or without ligands, were freshly prepared before each assay. For measurement of hydrolase activity, 3 mL reaction buffer containing enzyme, ε-NAD^+^, and ligands at indicated concentrations was maintained at 37 °C with constant stirring.

The cyclase activity of CD38 was measured as previously described (Higashida et al., 1999; Jin et al., 2007). CD38 cyclase activity was measured using 60 μM NGD as a substrate in cyclase reaction buffer (100 mM KCl, 10 μM CaCl_2_ and 50 mM Tris-HCl, pH 6.6). Tissues were cut into pieces and suspended in lysis buffer (5 mM MgCl_2_, 10 mM Tris-HCl, pH 7.3) at 4 °C for 30 min. Then, the suspension was homogenized on ice, and the supernatant was collected after centrifugation. To collect the crude membrane fractions, the supernatant was centrifuged at 105000 g for 30 min. The final pellet was resuspended in 10 mM Tris-HCl solution (pH 6.6) for cyclase assay. Homogenates of THP-1 or A549 cells, and recombinant hCD38 were used directly for cyclase assays. The loading volume of enzyme was 4 to 20 μL, the total volume of reaction system was 3 mL. To detect enzyme inhibition, except when specified, the ligands were directly dissolved in cyclase reaction buffer before pH adjustment, after which, the pH was immediately adjusted to 6.6. All reaction buffers, with or without ligands, were freshly prepared before each assay. For measurement of cyclase activity, 3 mL reaction buffer containing enzyme, NGD and ligands at indicated concentrations was maintained at 37 °C with constant stirring.

The reaction buffer for hydrolase or cyclase assays was excited at 300 nm, and fluorescence emission was measured every second at 410 nm by Shimazu RF-6000 spectrofluorometer. Hydrolase or cyclase activity was calculated from the linear portion of the time course by fitting a linear function to the data points recorded within 5 min.

In this study, 8-OG and guanosine were tested only at 50 μM due to the limited solubility. Guanosine was dissolved in DMSO and diluted in the reaction buffer for subsequent assays. Other ligands such as sUA, inosine, hypoxanthine, xanthine, allantoin, adenosine, uracil, 1,3-DHI-2-one, oxypurinol, caffeine, 1-MU, 1,3-DMU, 1,7-DMU, and 1,3,7-TMU were tested at the concentrations as indicated.

For reversibility test, recombinant hCD38 was pre-incubated for 30 min in 4 conditions: (1) control reaction buffer; (2) in the presence of 100 μM substrate (ε-NAD^+^ or NGD); (3) 500 μM sUA; (4) both substrate and sUA. Subsequently, the enzyme was diluted 100-fold in reaction buffer in the presence or absence of 500 μM sUA for activity assays using ε-NAD^+^ or NGD. Naïve THP-1 and A549 cells were incubated with RPMI-1640 medium in the presence or absence of 500 μM sUA for 2 h. The cells were collected and homogenized in the reaction buffer on ice with or without 500 μM sUA, samples were then diluted 100-fold in reaction buffer with or without 500 μM sUA for enzyme assay. Control group was not treated with sUA in all steps; sUA group was treated with sUA in each step; sUA release group was treated with sUA prior to dilution in sUA-free buffer for enzyme assays.

### Measurement of NAMPT and PARP activity

The activities of NAMPT and PARP were measured using commercial kits, HT Universal Colorimetric PARP Assay Kit and CycLex NAMPT Colorimetric Assay Kit Ver.2. The solutions of ligands were freshly prepared (pH was adjusted to 7.4) and were immediately used for enzyme assays.

### Moderate sUA supplementation in mice

Plasma sUA levels in mice were increased to the minimum physiological levels of humans by moderate sUA supplementation. For 1-day sUA supplementation, WT and CD38 KO mice received oral administration of saline, OA (1.5 g/kg), or OA (1.5 g/kg) plus inosine (1.5 g/kg), the gavage volume was 5 mL/kg. Drug suspension in saline was freshly prepared and warmed to 37 ℃ before each treatment. The mice received the first treatment on the evening (19:00) of the first day and the second treatment on the morning (9:00) of the second day. For 3-day or 7-day sUA supplementation, WT and CD38 KO mice received the same treatment twice daily from the evening of the first day to the morning of the last day. Four hours after the last treatment, the mice were sacrificed and whole blood, serum, plasma, and tissues were collected for metabolic studies. For immunological studies, mice were stimulated with different ligands 2 h after the second treatment on the morning (9:00) of the second day.

### sUA release in mice

WT mice received oral administration of saline, OA, or OA plus inosine (1-day supplementation model, as described above). One day (28 h) or 3 days (76 h) after the second treatment, the mice were sacrificed and whole blood and plasma were collected.

### cLPS-induced systemic inflammation

Plasma sUA levels in WT and CD38 KO mice were increased to the minimum physiological levels of humans by 1-day sUA supplementation (OA plus inosine). Two hours after the last treatment of OA, or OA plus inosine, the mice were intraperitoneally injected with sterile PBS or cLPS (2 mg/kg or 20 mg/kg), the injection volume was 3 mL/kg. Four hours (20 mg/kg) or 6 h (2 mg/kg) after stimulation, the mice were sacrificed and whole blood, plasma, and serum were collected.

### MSU crystal-induced peritonitis

Plasma sUA levels in WT and CD38 KO mice were increased to the minimum physiological levels of humans by 1-day sUA supplementation (OA plus inosine). Two hours after the last treatment of OA, or OA plus inosine, the mice were intraperitoneally injected with sterile PBS or MSU crystals (2 mg/mouse), and the injection volume was 200 μL/mouse. Six hours after stimulation, the mice were sacrificed and blood samples were collected. For each mouse, the peritoneal cavity was washed with 5 mL ice-cold sterile PBS, the supernatant was collected by centrifugation for subsequent ELISA, and cell pellets were used for total viable cell count by Trypan Blue staining. In brief, cell pellet from 1.5 mL of peritoneal lavage fluid was resuspended in RBC lysis buffer for 30 s, sterile PBS (9-fold volume) was added to terminate lysis. Afterward, the cells were centrifuged at 1000 rpm for 5 min and resuspended in sterile PBS for viable cell count. For neutrophil count, peritoneal lavage fluid was directly used for smears after appropriate dilution, and subsequently, Wright-Giemsa staining was performed according to the protocol provided by the manufacturer. Viable cells and neutrophils were counted by two investigators (one investigator did not participate in this project and was blind to the information of experimental groups), and the mean numbers were shown in the figures.

### ELISA

Human IL-1β, mouse IL-1β, IL-6, IL-18, TNF-α and CXCL1 levels were measured according to the protocols provided by the manufacturer. Serum samples were diluted before ELISA when applicable. In this study, all the samples were frozen after collection and thawed before ELISA.

### Preparation of MSU crystals and sUA solution

MSU crystals were prepared by the recrystallization of oversaturated sUA according to a previous report (Martinon et al., 2006). Improper preparation of sUA solution may introduce crystals to cause false-positive or false-negative results. It has been demonstrated that crystals may precipitate in sUA solution prepared by pre-warming to activate immune cells (Ma et al., 2020). In this study, the sUA solution was prepared according to an improved protocol (Ma et al., 2020). Briefly, we directly dissolved sUA at 0.5 mg/mL in blank RPMI-1640 medium by addition of NaOH, and adjusted the pH by HCl. Then sUA solution was immediately filtered by 0.2-μm filters and diluted to experimental concentration (up to 10 mg/dL, 595 μM). For all experiments, sUA solution was freshly prepared and used immediately.

### Preparation of BMDMs

Bone marrow cells were isolated from 10- to 12-week-old WT or CD38 KO mice (both male and female) by washing the marrow cavity with sterile PBS. The collected bone marrow cells were filtered by 70-μm strainers and then centrifuged at 1000 rpm, 4℃ for 5 min. The cell pellet was resuspended in 2 mL red blood cell (RBC) lysis buffer for 30 s, then 8 mL complete RPMI-1640 medium (10% FBS, 1% Penicillin/Streptomycin and 50 μM 2-mercaptoethanol) was added to terminate the lysis. After 5-min centrifugation at 1000 rpm, 4℃, the cells were resuspended in fresh complete RPMI-1640 medium and maintained for 4 h in an incubator. Adherent cells were discarded, whereas non-adherent cells were cultured in complete RPMI-1640 medium containing 20 ng/mL M-CSF. After 3-day differentiation, the medium was replaced with fresh complete RPMI-1640 medium containing 20 ng/mL M-CSF. On the 7^th^ day, BMDMs were collected for subsequent experiments. BMDMs were primed with 100 ng/mL ultrapure LPS for 4 h for canonical inflammasome assay. For metabolic assay of NMN, BMDMs were primed with 100 ng/mL ultrapure LPS for 8 h to induce higher protein expression of CD38.

### Intracellular NAD^+^ assay

A549 cells were seeded in 24-well plates for NAD^+^ assay. Briefly, when the confluence reached 80%, the culture medium in each well was discarded, and the cells were washed twice with sterile PBS and incubated in RPMI-1640 medium containing 1% FBS in the presence or absence of sUA (from 100 to 500 μM) for 20 h. Then the cells were washed twice with sterile PBS, after the second washing, PBS was completely removed and 100 μL 5% ice-cold PCA was added into each well. The plate was kept on ice for 2 h, then cell samples were collected and centrifuged at 15000 rpm, 4℃ for 10 min. The supernatant was used for subsequent handling and measurement (see LC-MS/MS analysis).

Naïve THP-1 cells were pre-incubated with sUA (0-10 mg/dL) in RPMI-1640 medium containing 1% FBS for 2h, then the cells were washed twice with sterile PBS and stimulated with MSU crystals, cLPS, zymosan or ATP for 6 h. Subsequently, the cells were washed twice with sterile PBS, and then a total of 100 μL 5% ice-cold PCA was used to extract NAD^+^ from cells in each well and medium as mentioned above.

WT BMDMs were primed with 100 ng/mL ultrapure LPS for 8 h. Then the cells were washed twice with sterile PBS. Afterward, the cells were incubated in control or 100 μM NMN-supplemented RPMI-1640 medium in the presence of sUA or 78c. After 6-h incubation, the cells were washed twice with sterile PBS. Finally, cell samples for NAD^+^ measurement were collected by 5% ice-cold PCA as mentioned above.

### Canonical inflammasome assay

Naïve THP-1 cells were primed with 0.5 μM phorbol 12-myristate 13-acetate (PMA) for 3 h the day before stimulation. Primed THP-1 cells were pre-incubated in RPMI-1640 medium in the presence or absence of sUA (5 or 10 mg/dL) for 2 h. The cells were then washed twice with sterile PBS, and were stimulated with MSU crystals, cLPS, zymosan, and ATP in serum-free RPMI-1640 medium for 4 h.

WT and CD38 KO BMDMs were primed with 100 ng/mL ultrapure LPS for 4 h. Subsequently, primed BMDMs were pre-incubated with or without sUA (5 or 10 mg/dL) for 2 h, the cells were then washed twice with sterile PBS and were stimulated with nigericin, MSU crystals, or cLPS in serum-free RPMI-1640 medium.

After stimulation, the culture medium was collected and centrifuged at 3000 rpm, 4 ℃ for 5 min, the supernatant was collected and stored at −30 ℃ until ELISA.

### sUA uptake assay

WT BMDMs were primed with 100 ng/mL ultrapure LPS for 4 h. Then the cells were washed twice with sterile PBS, and maintained in RPMI-1640 medium containing sUA (100, 200, or 500 μM) for 2 h or 15 h. Subsequently, the cells were washed twice with ice-cold sterile PBS to terminate uptake, 100 μL 5% ice-cold PCA was added into each well. The plates were placed on ice for 2 h, then cell samples were collected and centrifuged at 15000 rpm, 4℃ for 10 min. The supernatant was used for subsequent handling and measurement (see LC-MS/MS analysis).

### Metabolic assay of NAD^+^

NAD^+^ degradation by recombinant hCD38 was detected in hydrolase reaction buffer. At first, sUA (100, 200 and 500 μM) or other ligands at 500 μM in hydrolase reaction buffer (250 mM sucrose, 40 mM Tris) was freshly prepared, then the pH was immediately adjusted to 7.4 by HCl. Recombinant hCD38 was added into the buffer of experimental groups at 20 ng/mL, then the buffer was maintained at 37 ℃. The substrate NAD^+^ was dissolved in hydrolase reaction buffer (pH 7.4) at 10 mM, and the pH was further adjusted to 7.4. The reaction was started with the addition of NAD^+^ in the buffer for each group (final concentration is 200 μM), including control group (recombinant hCD38 free). All the buffers were incubated at 37 ℃ with constant stirring. After 30 or 60 min, 20 μL reaction buffer was transferred into 180 μL of 5% ice-cold PCA, then vortexed for 30 s before 10-min centrifugation at 15000 rpm, 4 ℃. The supernatant was further handled for NAD^+^ measurement within 12 h without freezing and thawing (see LC-MS/MS analysis).

### Metabolic assay of NMN

To prepare the reaction buffer, sUA was directly dissolved in blank RPMI-1640 medium by addition of NaOH, and the pH was immediately adjusted by HCl. After pH adjustment, the medium containing sUA was filtered by 0.2-μm filter and diluted as indicated. The medium containing 20 ng/mL recombinant hCD38 was then placed in a cell incubator. The reaction was started with the addition of NMN (final concentration was 200 μM). After incubation for 6 h, 20 μL medium was transferred into 180 μL 5% ice-cold PCA, then vortexed for 30 s and centrifuged at 15000 rpm, 4 ℃ for 10 min. The supernatant was stored at −80 ℃ until further handling for NMN measurement (see LC-MS/MS analysis).

### Extracellular NMN degradation

WT and CD38 KO BMDMs were primed with 100 ng/mL ultrapure LPS for 8 h, then the cells were washed twice with sterile PBS. Primed BMDMs were maintained in RPMI-1640 medium supplemented with 100 μM NMN in the presence or absence of sUA. To restore the inhibitory effects of sUA on NMN degradation in KO BMDMs, recombinant hCD38 was added to the medium (final concentration was 10 ng/mL). After 6-h incubation, culture medium was collected and centrifuged at 3000 rpm, 4 ℃ for 5 min. Then, 20 μL of supernatant was transferred into 180 μL 5% ice-cold PCA, and was vortexed for 30 s and centrifuged at 15000 rpm, 4 ℃ for 10 min. The supernatant was stored at −80 ℃ until further handling for NMN measurement.

### Handling of animal samples

The collected whole blood samples were immediately diluted 10-fold in 5% ice-cold perchloric acid (PCA) and homogenized on ice. After 10-min centrifugation at 15000 rpm, 4 ℃, the supernatant was subpackaged for measurement or −80 ℃ storage. For NAD^+^ measurement in whole blood, the supernatant was handled without freezing and thawing and was measured within 24 h of sample collection. For measurement of plasma sUA, after sample collection, we immediately diluted the plasma with 5% ice-cold PCA, after vortex, samples were centrifuged at 15000 rpm, 4 ℃ for 10 min, the collected supernatant was used for the subsequent handling and measurement within 24 h.

After the collection of blood samples, the mice were immediately perfused with ice-cold sterile PBS. The tissue samples were collected, dried with tissue paper, and weighed. All the tissue samples were immediately homogenized in 5% ice-cold PCA on ice, then centrifuged at 15000 rpm, 4℃ for 10 min. The supernatant was collected and stored at −80 ℃ until further handling for measurement (see LC-MS/MS analysis).

### LC-MS/MS analysis

As mentioned above, samples (cells, tissues, blood or reaction buffer) were treated with ice-cold PCA, after extraction and centrifugation, the collected supernatant was appropriately diluted in 5% ice-cold PCA when applicable, and then was vortexed for 30 s and centrifuged at 15000 rpm, 4℃ for 5 min. Afterward, 30 μL supernatant was added into 200 μL 5 mM ammonium formate containing internal standards at indicated concentrations. Then all the samples were vortexed for 30 s and centrifuged at 15000 rpm, 4℃ for 5 min again, the final supernatant was used for subsequent measurement. All the samples after handling were immediately analyzed in this study without freezing and thawing.

The LC-MS/MS system consisted of a triple quadrupole LCMS-8050 (Shimadzu) and an LC-30A system (Shimadzu). NAD^+^, NMN and the N-cyclohexyl benzamide (NCB, internal standard, 5 ng/mL) were eluted on Altlantis® HILIC Silica (2.1 × 150 mm, 5 μm) at 40 °C using an isocratic mobile phase containing 60% water with 0.1% formic acid and 40% acetonitrile with 0.1% formic acid at 0.4 mL/min. The selected transitions of m/z were 664.10 → 136.10 for NAD^+^, 334.95 → 123.15 for NMN, and 204.10 → 122.20 for NCB in positive ion mode. NAD^+^ and NMN were measured independently in this study, as their peaks were hardly separated within a short time.

sUA, cADPR and FYU-981 (internal standard, 1 μM, Fuji Yakuhin Co., Ltd.) were eluted on CAPCELL PAK C8 TYPE UG 120 (2 × 150 mm, 5 μm) at 40 °C. sUA and FYU-981 were separated using a gradient mobile phase containing water with 0.1% formic acid (A) and acetonitrile with 0.1% formic acid (B) at 0.4 mL/min. The elution was started with 55% A for 0.5 min, A was decreased from 55% to 5% for 1 min, then A was increased from 5% to 55% for 0.5 min, finally 55% A was maintained for 0.5 min. cADPR and FYU-981 were separated using an isocratic mobile phase containing 60% water with 0.1% formic acid and 40% acetonitrile with 0.1% formic acid at 0.4 mL/min. The selected transitions of m/z were 167.10 → 124.10 for sUA, 539.95 → 272.95 for cADPR, and 355.95 → 159.90 for FYU-981 in negative ion mode.

The injection volume was 1 μL for all measurements, and data manipulation was accomplished by Labsolutions software (version 5.97, Shimadzu).

### Statistical analysis

Statistics were performed using GraphPad Prism 9. Sample size was not predetermined by any statistical methods. Comparisons between multiple groups were performed using 1-way ANOVA with Dunnett’s or Tukey’s multiple comparisons test, Kruskal-Wallis test, Brown-Forsythe and Welch ANOVA tests, or 2-way ANOVA with Tukey’s multiple comparisons test. Two-tailed unpaired t-test (with Welch’s correction when applicable), or Mann-Whitney test were used for the analysis between two groups when applicable. Data were shown as mean ± s.e.m. or mean ± s.d. as indicated. *P* < 0.05 was considered significant. The details of statistical tests and the numbers of mice or biologically independent samples were described in the figure legends.

## Supporting information

Key Resources Table

## Author contributions

Shijie Wen: Conceptualization, Data curation, Formal analysis, Methodology, Investigation, Validation, Visualization, Writing-original draft, Writing-review & editing, Project administration.

Hiroshi Arakawa: Data curation, Formal analysis, Methodology, Writing-review & editing, Funding acquisition, Project administration.

Shigeru Yokoyama: Resources, Writing-review & editing.

Yoshiyuki Shirasaka: Resources, Writing-review & editing.

Haruhiro Higashida: Methodology, Resources, Writing-review & editing.

Ikumi Tamai: Conceptualization, Data curation, Formal analysis, Funding acquisition, Writing-review & editing, Project administration, Supervision.

## Acknowledgments

This study was supported by KAKENHI [JP21H02641] (I.T.), [JP23K18181] (I.T.), and [JP22K19372] (H.A.) from the Japan Society for the Promotion of Science (JSPS) and Research Grant 2022 (I.T.) from Gout and Uric Acid Foundation of Japan. S.W. was funded by the Japanese Government (Monbukagakusho: MEXT) Scholarship Program and Kanazawa University. The authors thank Dr. Zheng Jing, Ms. Aimi Taniguchi, Dr. Anpei Zhang, Mr. Kazuki Himi, and Mr. Kazuki Fujita for providing assistance.

## Data availability

All data generated or analyzed in this study are provided in the manuscript and supporting information.

## Figure supplements

**Figure 1-figure supplement 1.**
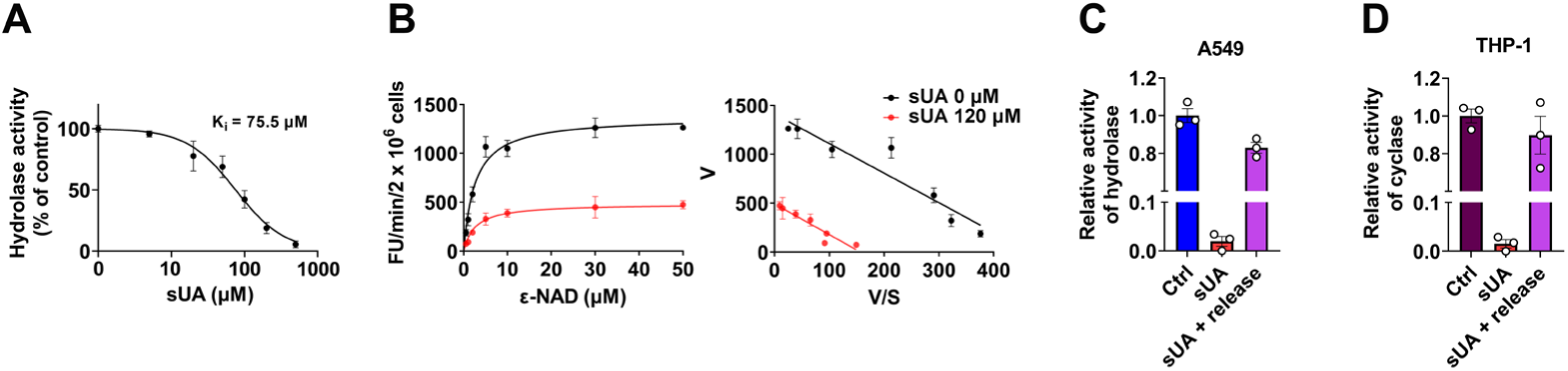
sUA inhibition of CD38 and reversibility in THP-1 and A549 cells. **(A)** Hydrolase activity of THP-1 cells in the presence of sUA (0 to 500 μM) (n = 3 experiments/technical replicates). **(B)** Effect of different ε-NAD^+^ concentrations on sUA inhibition of hydrolase activity of THP-1 cells (n = 3 experiments/technical replicates). **(C** and **D)** Reversibility of inhibition of hydrolase (A549 cells) (**C**) and cyclase (THP-1 cells) (**D**) by sUA (n = 3 experiments/technical replicates). Data are mean ± s.d. (**A** and **B**) or mean ± s.e.m. (**C** and **D**).

**Figure 1-figure supplement 2.**
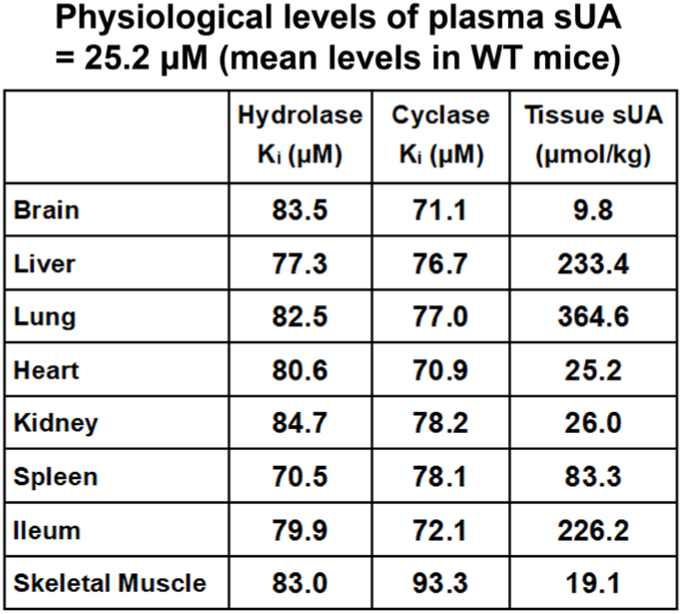
Comparison between K_i_ values and mean levels of sUA in different tissues. K_i_ values were also shown in Figure 1E, and serum and tissue sUA levels were from WT mice that received 1-day treatment of saline (also shown in Figure 3D and Figure 3-figure supplement 5A).

**Figure 1-figure supplement 3.**
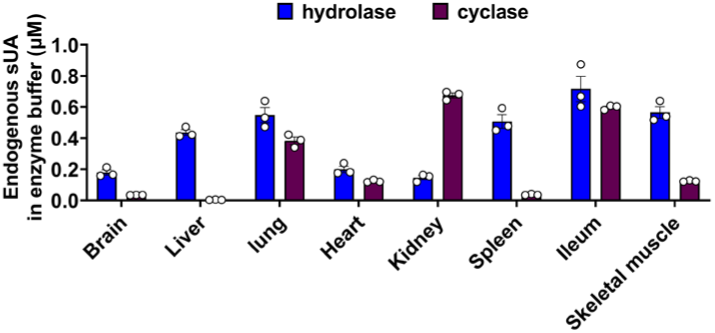
Endogenous sUA concentrations in the final reaction buffer for enzyme assays. sUA levels in initial homogenate or membrane fractions were measured, then the endogenous sUA concentrations in the final reaction buffer were calculated based on loading dilution (n = 3 biologically independent samples). Data are mean ± s.e.m.

**Figure 2-figure supplement 1.**
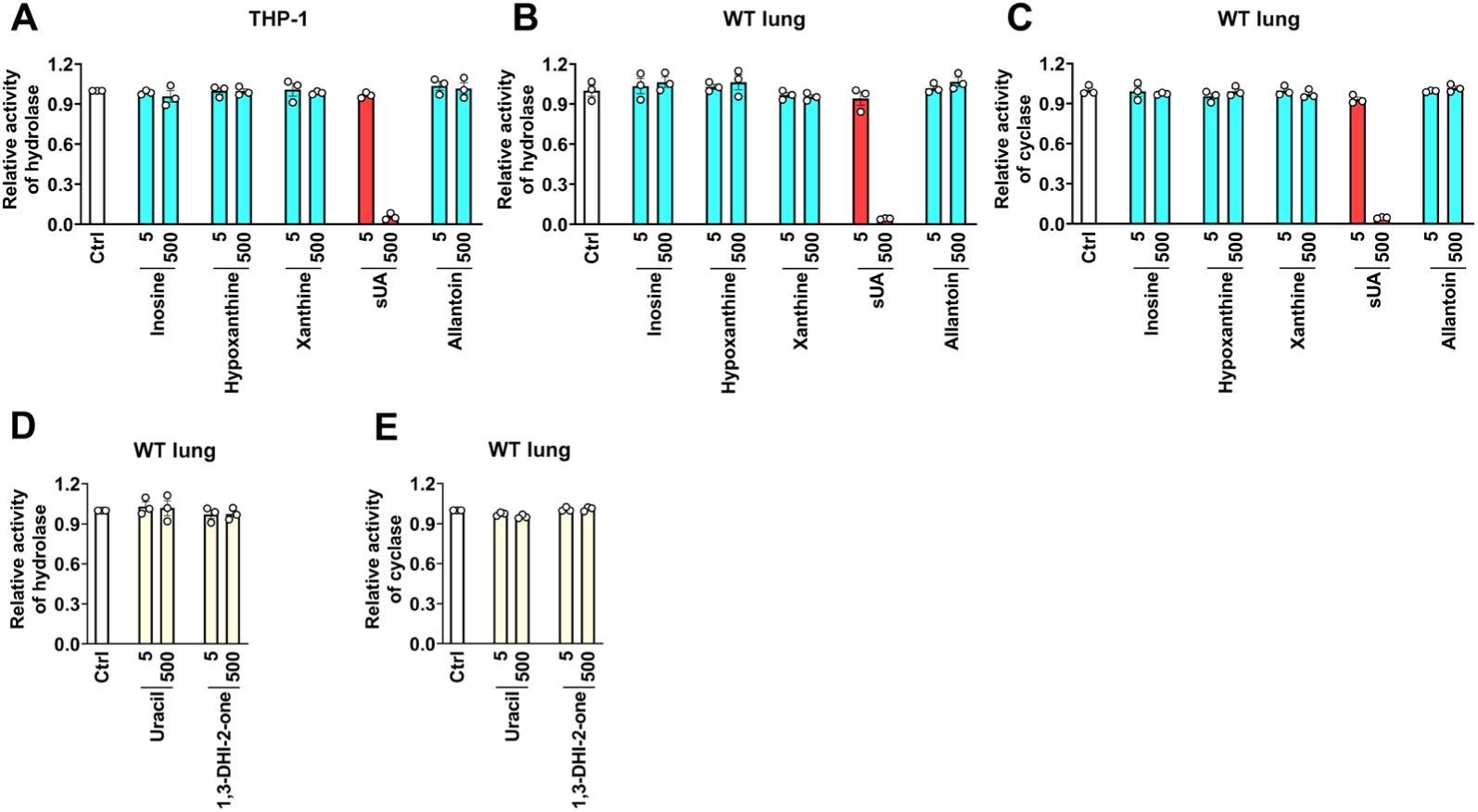
Effect of sUA, sUA precursors and metabolite, and other derivates on CD38 activity in cells and tissues. **(A-C)** Effect of sUA and its precursors and metabolite on hydrolase (**A** and **B**) and cyclase (**C**) activities. (n = 3 experiments/technical replicates) **(D** and **E)** Effect of uracil and 1,3-dihydroimidazol-2-one (1,3-DHI-2-one) on hydrolase and cyclase activities of WT lung tissues. (n = 3 experiments/technical replicates) Data are mean ± s.e.m.

**Figure 3-figure supplement 1.**
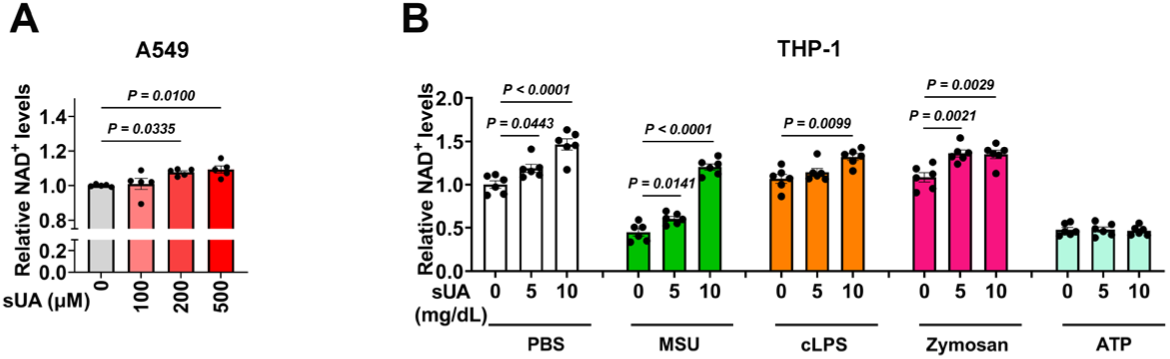
Effect of sUA on intracellular NAD^+^ levels in A549 and THP-1 cells. **(A)** Effect of sUA on intracellular NAD^+^ levels of A549 cells (n = 5 biologically independent samples). **(B)** Effect of sUA pre-incubation on intracellular NAD^+^ levels of THP-1 cells. Naïve THP-1 cells were incubated with sUA (0-10 mg/dL) for 2h, then the cells were washed twice with sterile PBS and stimulated with MSU crystals (200 μg/mL), cLPS (20 μg/mL), zymosan (50 μg/mL) or ATP (2 mM) for 6 h (n = 6 biologically independent samples). Data are mean ± s.e.m. Significance was tested using 1-way ANOVA with Dunnett’s multiple comparisons test (**A** and **B**) or Kruskal-Wallis test (cLPS groups in **B**).

**Figure 3-figure supplement 2.**
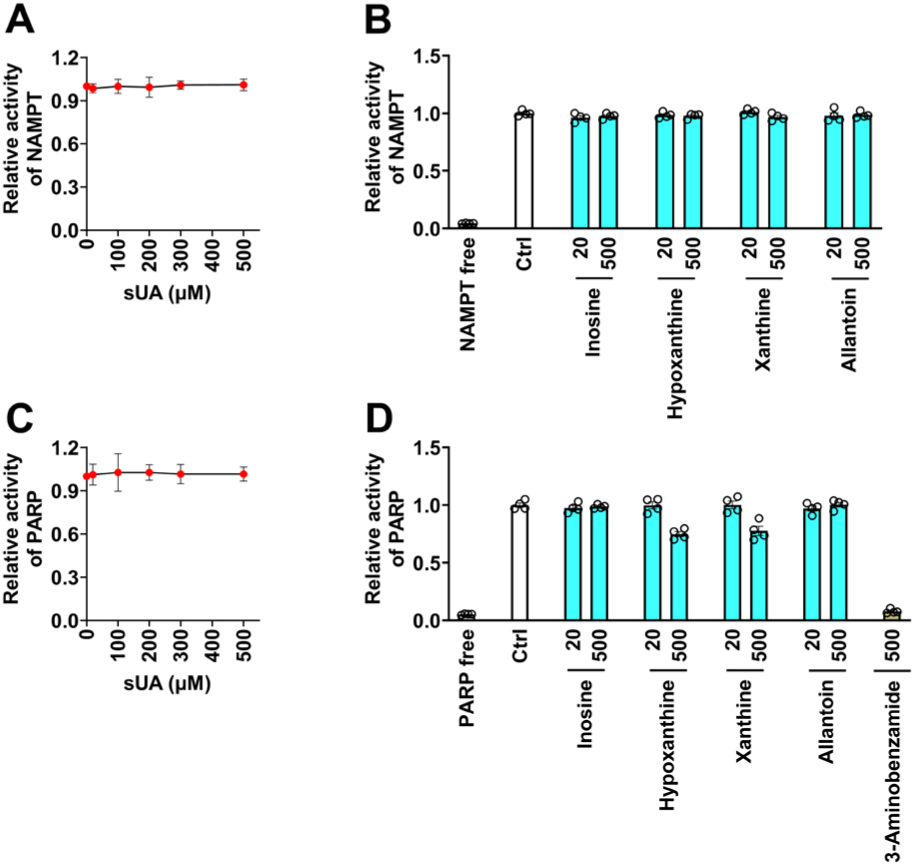
Effect of sUA and its precursors and metabolite on NAMPT and PARP activities. **(A and B)** NAMPT activity. (n = 4 experiments/technical replicates) **(C** and **D)** PARP activity. 3-Aminobenzamide is a PARP inhibitor. (n = 4 experiments/technical replicates) Data are mean ± s.d. (**A** and **C**) or mean ± s.e.m. (**B** and **D**).

**Figure 3-figure supplement 3.**
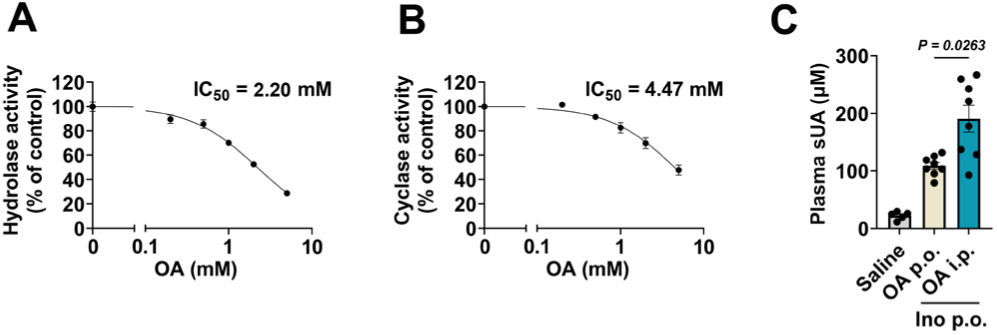
Effect of OA on CD38 activity and plasma sUA levels. **(A and B)** Hydrolase and cyclase activities of lung tissues from WT mice in the presence of OA (0 to 5 mM) (n = 3 experiments/technical replicates). **(C)** Effect of OA administration on plasma sUA levels in WT mice (10- to 12-week-old) that received oral administration of inosine. In saline group, the mice received oral administration and intraperitoneal injection of saline. In OA p.o. group, the mice received oral administration of inosine (1.5 g/kg) and OA (1.5 g/kg) (the same treatment in our models), and intraperitoneal injection of saline. In OA i.p. group, the mice received oral administration of inosine (1.5 g/kg), and intraperitoneal injection of OA (0.25 g/kg). Four hours after treatment, plasma sUA was measured (n = 5-8 mice per group). Data are mean ± s.d. (**A** and **B**) or mean ± s.e.m. (**C**). Significance was tested using Brown-Forsythe and Welch ANOVA tests (**C**).

**Figure 3-figure supplement 4.**
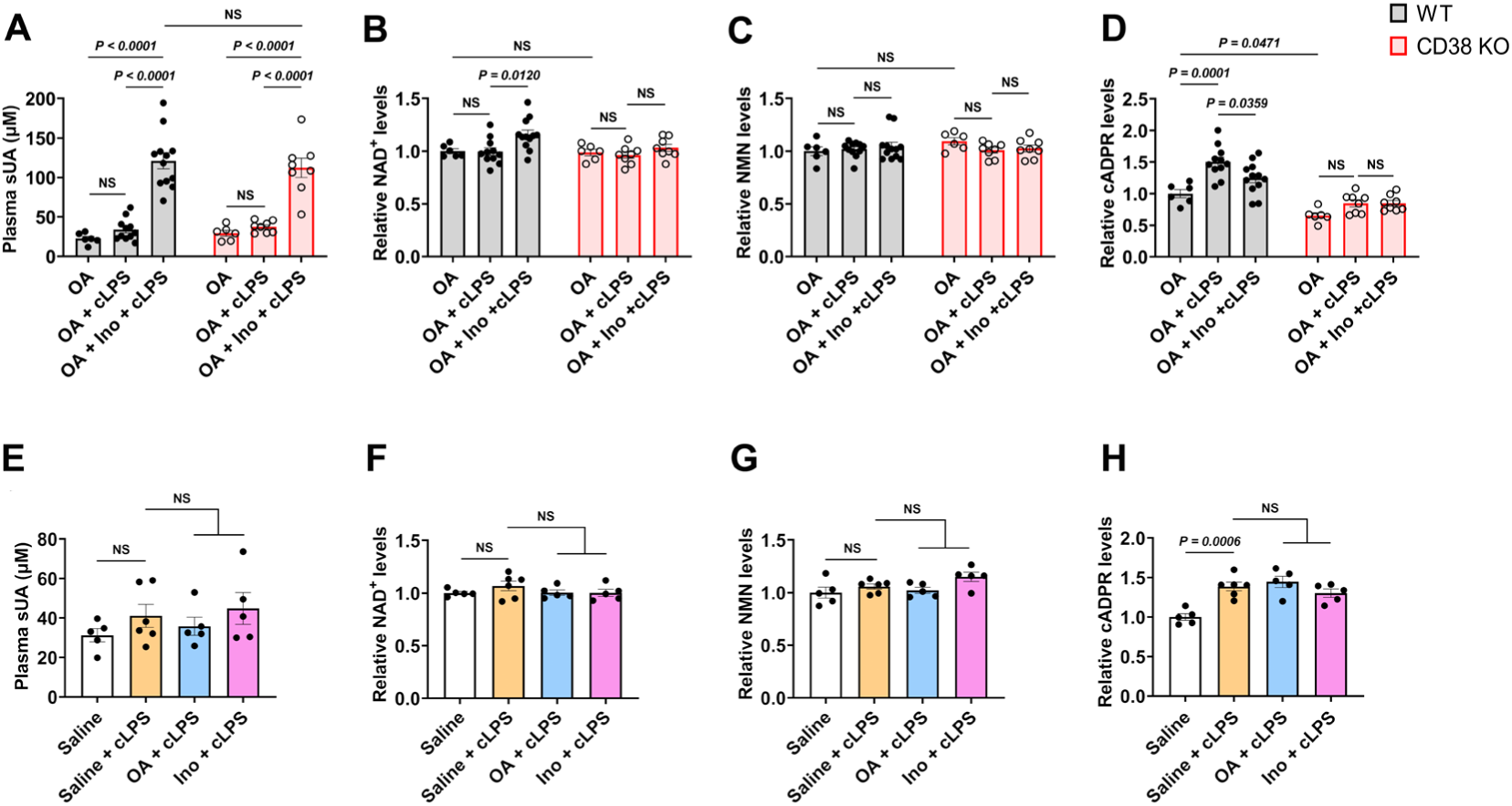
Effect of sUA at physiological levels on NAD^+^ degradation under inflammatory conditions. **(A-D)** Effect of 1-day sUA supplementation on plasma sUA (**A**) and whole blood NAD^+^(**B**), NMN (**C**), and cADPR (**D**) levels in WT and CD38 KO mice (10- to 12-week-old) under inflammatory conditions. The mice received 1-day sUA supplementation. Two hours after the last treatment, the mice were intraperitoneally stimulated with sterile PBS or cLPS (2 mg/kg) for 6 h (WT-OA: n = 6 mice, WT-OA + cLPS: n = 11 mice, WT-OA + Ino + cLPS: n = 12 mice, KO-OA: n = 6 mice, KO-OA + cLPS: n = 8 mice, KO-OA + Ino + cLPS: n = 8 mice). **(E-H)** Effect of 1-day treatment of OA or inosine (Ino) on plasma sUA (**E**) and whole blood NAD^+^ (**F**), NMN (**G**), and cADPR (**H**) levels in WT mice (10- to 12-week-old) under inflammatory conditions. The mice received 1-day treatment of OA or Ino (from the evening of day 0 to the morning of day 1). Two hours after the last treatment, the mice were intraperitoneally stimulated with sterile PBS or cLPS (2 mg/kg) for 6 h (n = 6 mice in Saline + cLPS group, n = 5 mice in other groups). Data are mean ± s.e.m. Significance was tested using 2-way ANOVA with Tukey’s multiple comparisons test (**A-D**), or 1-way ANOVA with Tukey’s multiple comparisons test (**E-H**). Statistic difference (**A**-**D**) between OA + cLPS and OA + Ino + cLPS groups in WT or KO mice (OA alone group excluded) was also analyzed by unpaired two-sided t-test (with Welch’s correction when applicable) or Mann Whitney test; WT mice: *P* < 0.0001 (**A**), or *P* = 0.0092 (**B**), 0.0221 (**D**); KO mice: *P* = 0.0004 (**A**).

**Figure 3-figure supplement 5.**
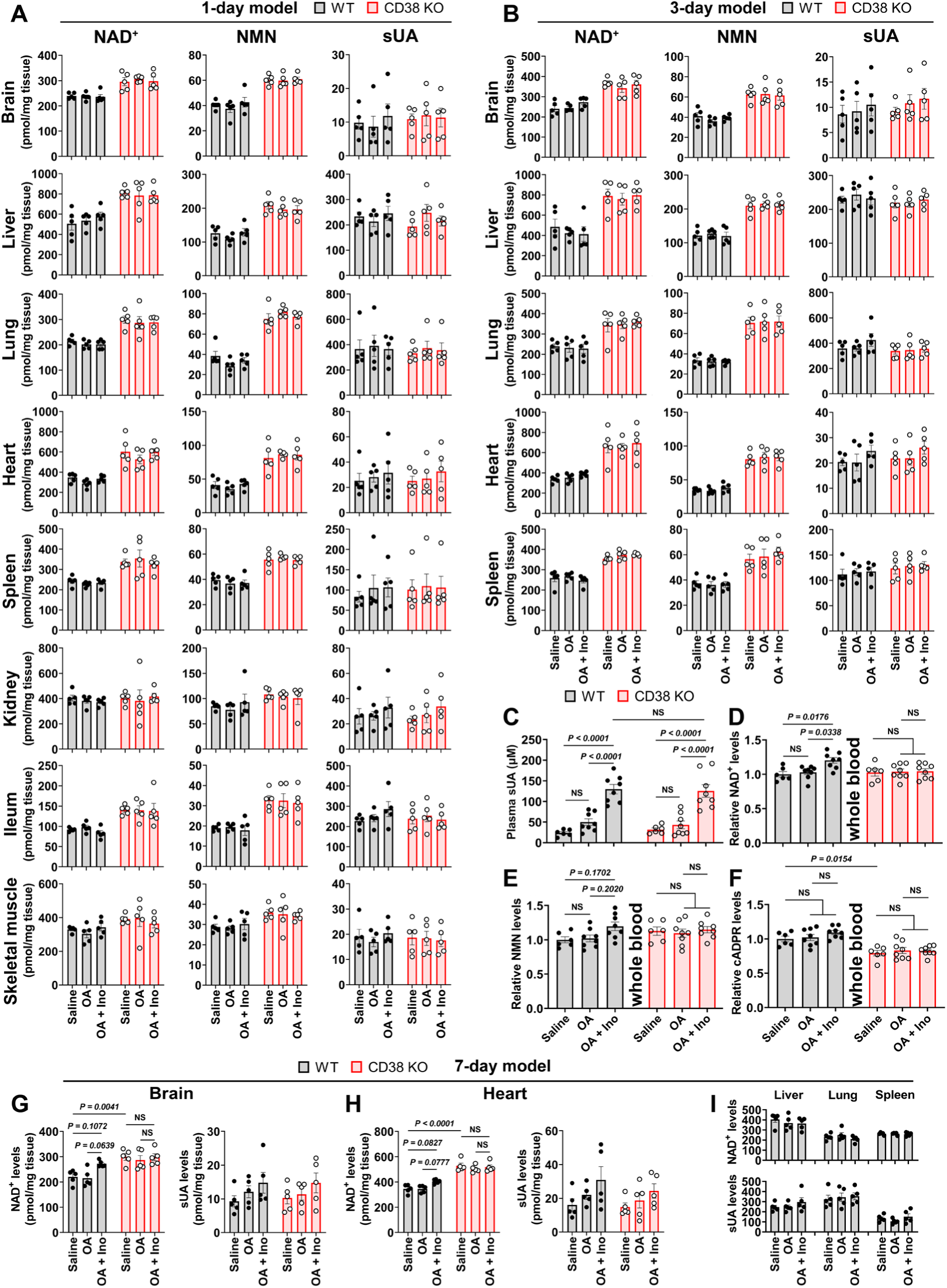
Effect of 1-day, 3-day, and 7-day moderate sUA supplementation on NAD^+^, NMN, and sUA levels in whole blood and tissues. WT and CD38 KO mice (10- to 12-week-old) received oral administration of saline, OA, or OA plus inosine (Ino) twice daily for 1, 3, or 7 days. Four hours after the last treatment, the mice were sacrificed. In 1-day model, the mice were treated from the evening of day 0 to the morning of day 1. **(A)** Effect of 1-day sUA supplementation on tissue NAD^+^, NMN, and sUA levels (n = 5 male mice per group). **(B)** Effect of 3-day sUA supplementation on tissue NAD^+^, NMN, and sUA levels (n = 5 male mice per group). **(C-F)** Effect of 3-day sUA supplementation on plasma sUA (**C**), whole blood NAD^+^ (**D**), NMN (**E**), and cADPR (**F**) levels (WT-Saline: n = 6 mice, WT-OA: n = 8 mice, WT-OA + Ino: n = 8 mice, KO-Saline: n = 6 mice, KO-OA: n = 8 mice, KO-OA + Ino: n = 8 mice). **(G-I)** Effect of 7-day sUA supplementation on tissue NAD^+^ and sUA levels (n = 5 male mice per group). Data are mean ± s.e.m. Significance was tested using 2-way ANOVA with Tukey’s multiple comparisons test (**A-H**), Kruskal-Wallis test or 1-way ANOVA with Dunnett’s multiple comparisons test (**I**). Statistic difference (**C**-**H**) between OA and OA + Ino groups in WT or KO mice (saline alone group excluded) was also analyzed by unpaired two-sided t-test; WT mice: *P* < 0.0001 (**C**), or *P* = 0.0058 (**D**), 0.0453 (**E**), 0.0083 (**G,** left panel), 0.0036 (**H,** left panel); KO mice: *P* = 0.0004 (**C**).

**Figure 3-figure supplement 6.**
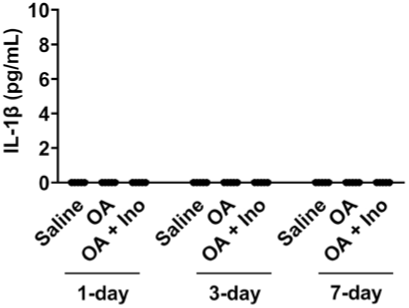
No effect of 1- to 7-day sUA supplementation on serum IL-1β production. Four hours after the last treatment, serum was collected for IL-1β measurement (n = 5 mice per group).

**Figure 3-figure supplement 7.**
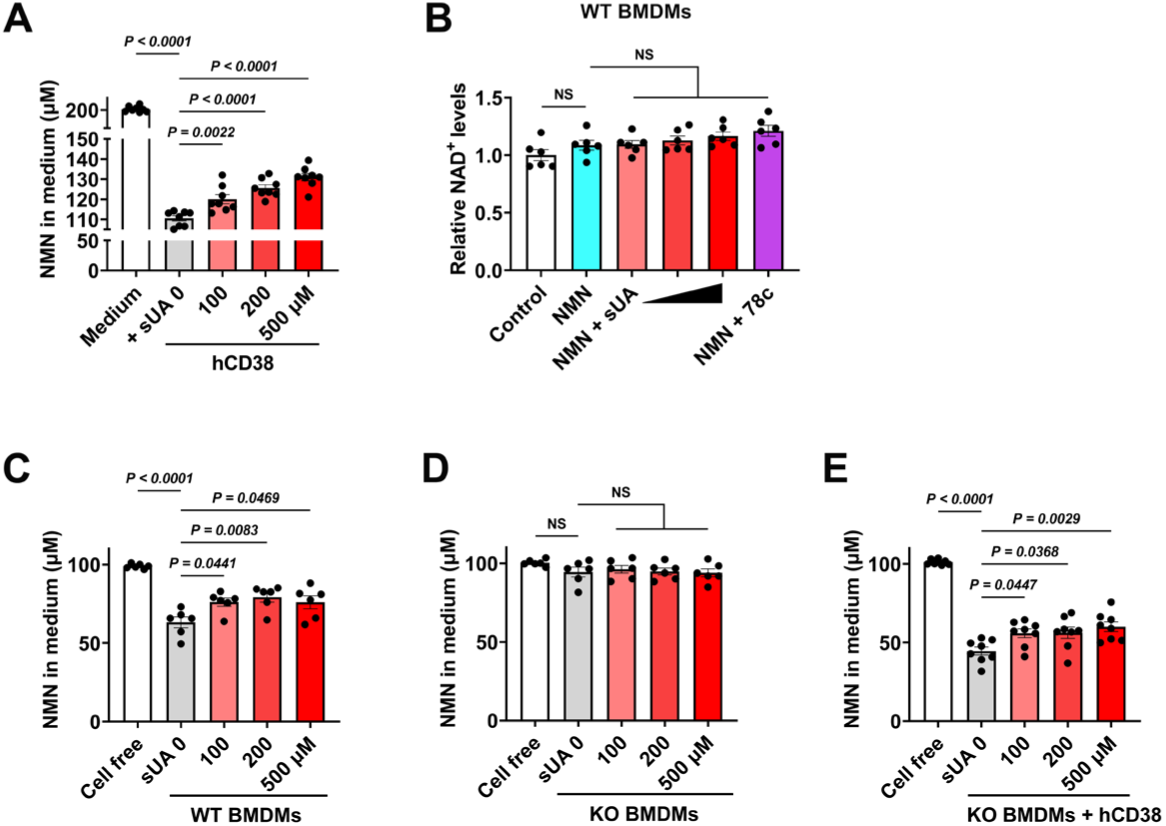
sUA limits NMN degradation via CD38 inhibition. **(A)** Effect of sUA on recombinant hCD38-mediated NMN (200 μM) degradation in medium (n = 8 independent samples). **(B)** Effect of sUA (100, 200, and 500 μM) or 78c (0.5 μM) on intracellular NAD^+^ levels of WT BMDMs treated with NMN. WT BMDMs were primed with100 ng/mL ultrapure LPS for 8 h (n = 6 biologically independent samples). **(C-E)** Effect of sUA on extracellular NMN degradation in WT (**C**) or CD38 KO BMDMs in the absence (**D**) or presence (**E**) of recombinant hCD38 (10 ng/mL). BMDMs were primed with 100 ng/mL ultrapure LPS for 8 h before metabolic assays. (n = 6 biologically independent samples in **C** and **D**, n = 8 biologically independent samples in **E**) Data are mean ± s.e.m. Significance was tested using 1-way ANOVA with Tukey’s multiple comparisons test.

**Figure 4-figure supplement 1.**
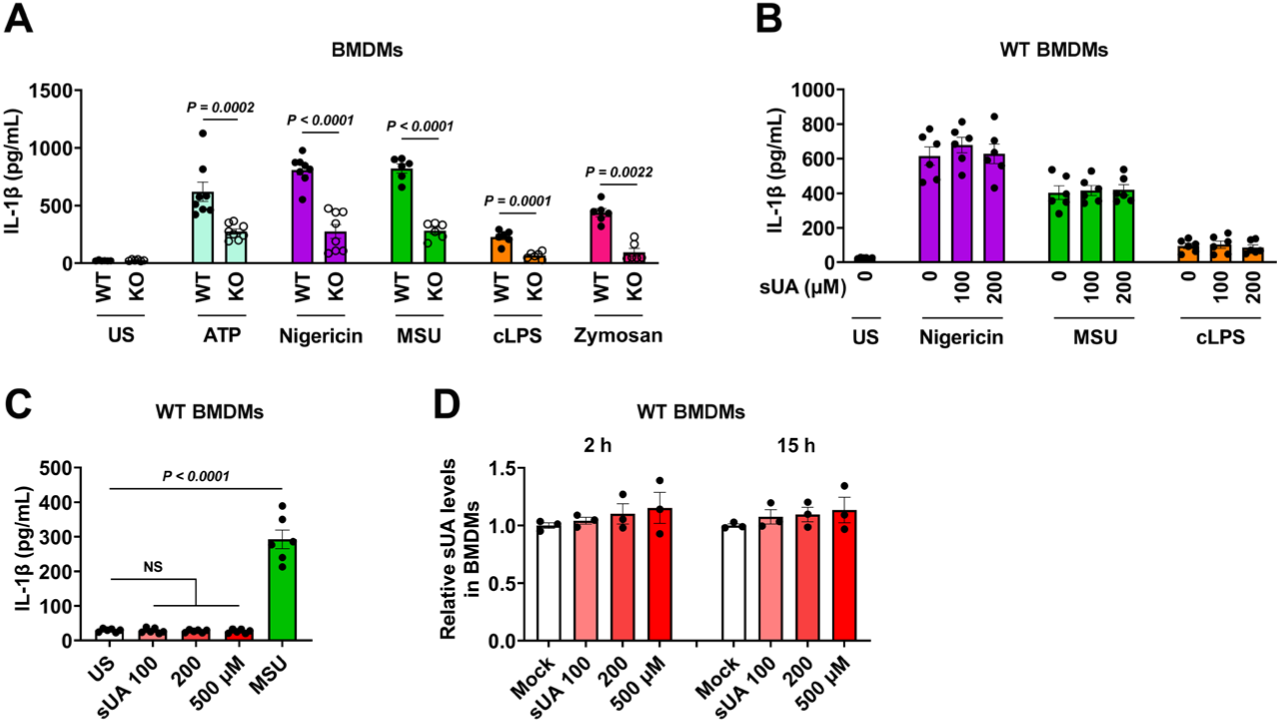
Effect of CD38 KO and sUA pre-incubation on IL-1β release in primed BMDMs. **(A)** Effect of CD38 KO on IL-1β release in primed BMDMs. WT and KO BMDMs were primed with 100 ng/mL ultrapure LPS for 4 h, then primed BMDMs were challenged by ATP (5 mM, 30min), nigericin (3 μM, 2 h), MSU crystals (200 μg/mL, 6 h), cLPS (20 μg/mL, 6 h), and zymosan (50 μg/mL, 4 h). US means unstimulated. (n = 8 biologically independent samples in ATP and nigericin groups, n = 6 biologically independent samples in other groups) **(B-D)** Effect of sUA pre-incubation on IL-1β release and intracellular sUA levels in primed BMDMs. WT BMDMs were primed with 100 ng/mL ultrapure LPS for 4 h. **(B)** The cells were pre-incubated with or without sUA (100 or 200 μM) for 2 h. Then, the cells were washed twice with sterile PBS and were stimulated with nigericin (3 μM, 2 h), MSU crystals (200 μg/mL, 4 h), or cLPS (1 μg/mL, 4 h). (**C** and **D)** Primed BMDMs were directly incubated with sUA (100, 200, and 500 μM) or MSU crystals (100 μg/mL) for 6 h in **C**, 2 or 15 h in **D**. US means unstimulated. (n = 6 biologically independent samples in **B** and **C,** n = 3 biologically independent samples in **D**) Data are mean ± s.e.m. Significance was tested using two-tailed unpaired t-test and Mann-Whitney test (ATP and Zymosan) (**A**), or 1-way ANOVA with Dunnett’s multiple comparisons test (**B-D**).

**Figure 4-figure supplement 2.**
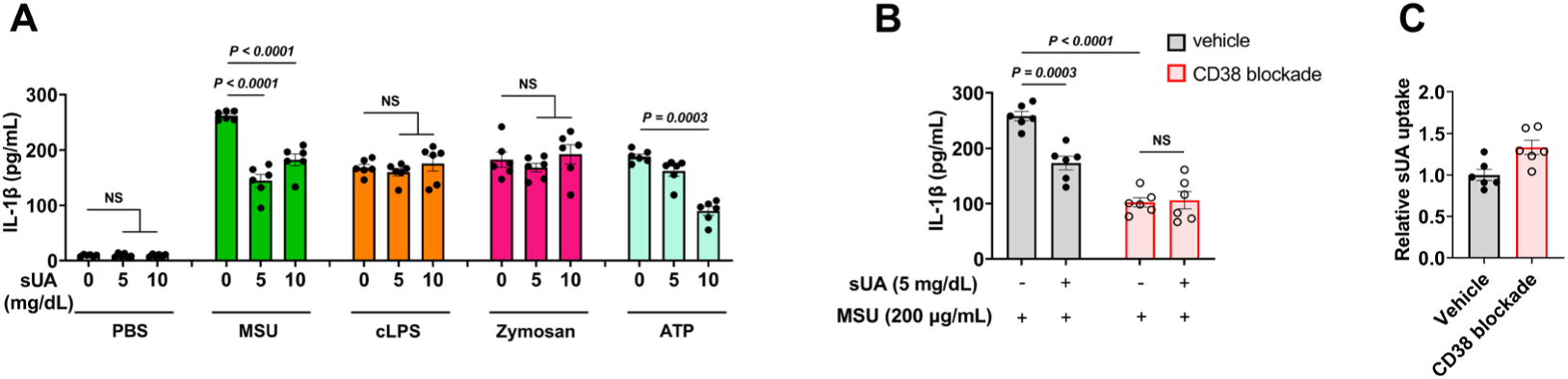
Effect of sUA pre-incubation and CD38 blockade on IL-1β release in primed THP-1 cells. THP-1 cells were primed with 0.5 μM PMA for 3 h the day before stimulation. (**A**) Primed THP-1 cells were pre-incubated with sUA (0-10 mg/dL) for 2 h, then the cells were washed twice with sterile PBS and challenged by MSU crystals (200 μg/mL), cLPS (20 μg/mL), zymosan (50 μg/mL), and ATP (2 mM) for 4 h (n = 6 biologically independent samples). **(B** and **C)** Primed THP-1 cells were treated without or with 2 μM 78c (CD38 blockade by the highly specific and potent inhibitor of CD38) for 2 h. Then the cells were pre-incubated with blank medium or sUA for 2 h. The cells were washed twice with sterile PBS. The washed cells were challenged by MSU crystals (200 μg/mL, 4 h) for IL-1β measurement (**B**) or were directly used for sUA uptake measurement without MSU crystal stimulation (**C**). 78c was used in all steps for CD38 blockade groups, including sUA pre-incubation and MSU crystal stimulation (n = 6 biologically independent samples). Data are mean ± s.e.m. Significance was tested using Kruskal-Wallis test (ATP groups in A), 1-way ANOVA with Dunnett’s multiple comparisons test (**A)**, or 2-way ANOVA with Tukey’s multiple comparisons test (**B**).

**Figure 4-figure supplement 3.**
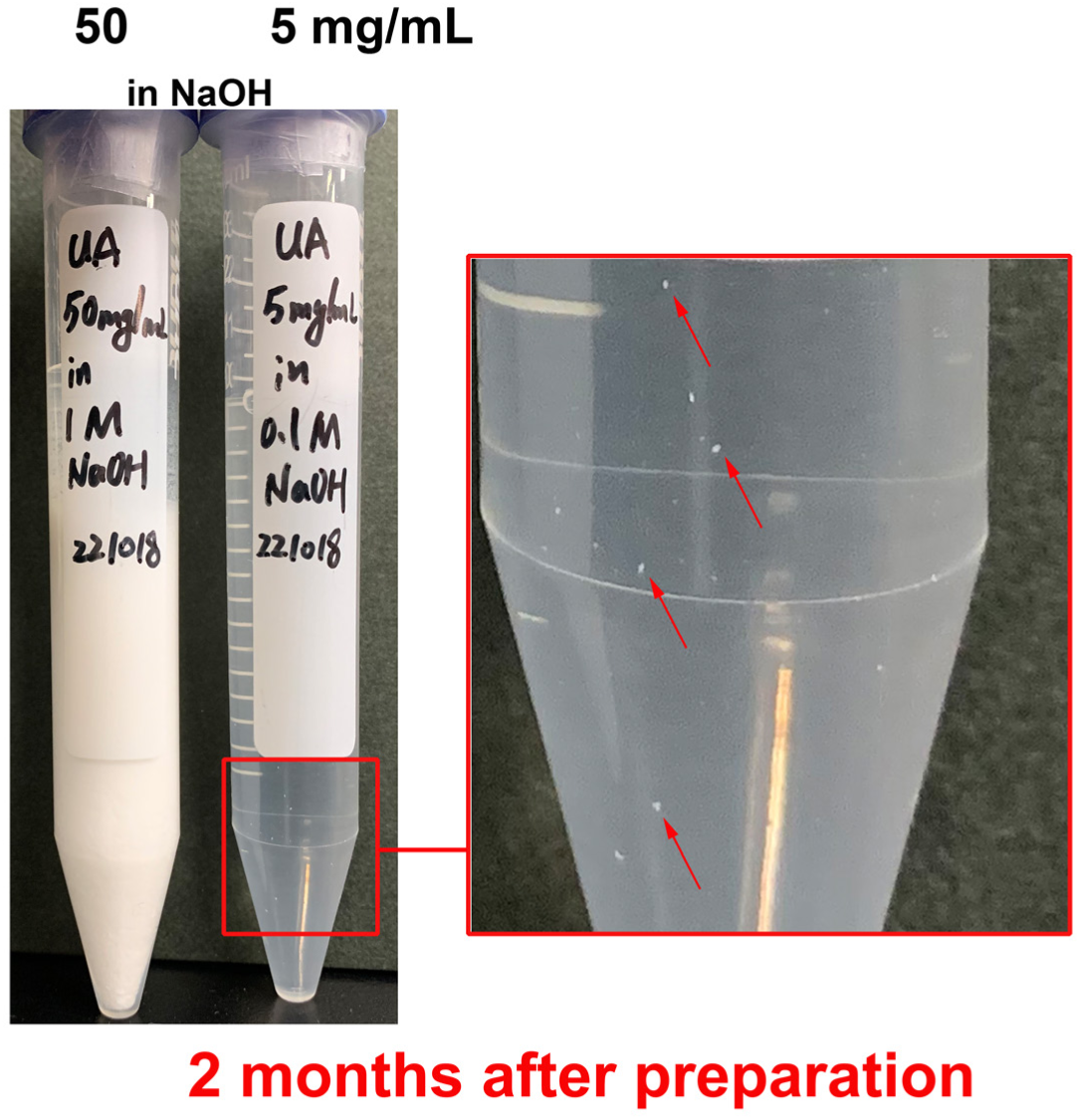
Crystal precipitation in high-concentration sUA stock solutions after long-term storage. sUA stock solutions in NaOH were prepared without pH adjustment. Crystals were immediately precipitated after dissolution in 50 mg/mL tube. Visible crystals were observed in 5 mg/mL tube after 2-month storage at 4 ℃ (before taking the picture, the tube was slightly shaken to resuspend the crystals on the bottom).

**Figure 4-figure supplement 4.**
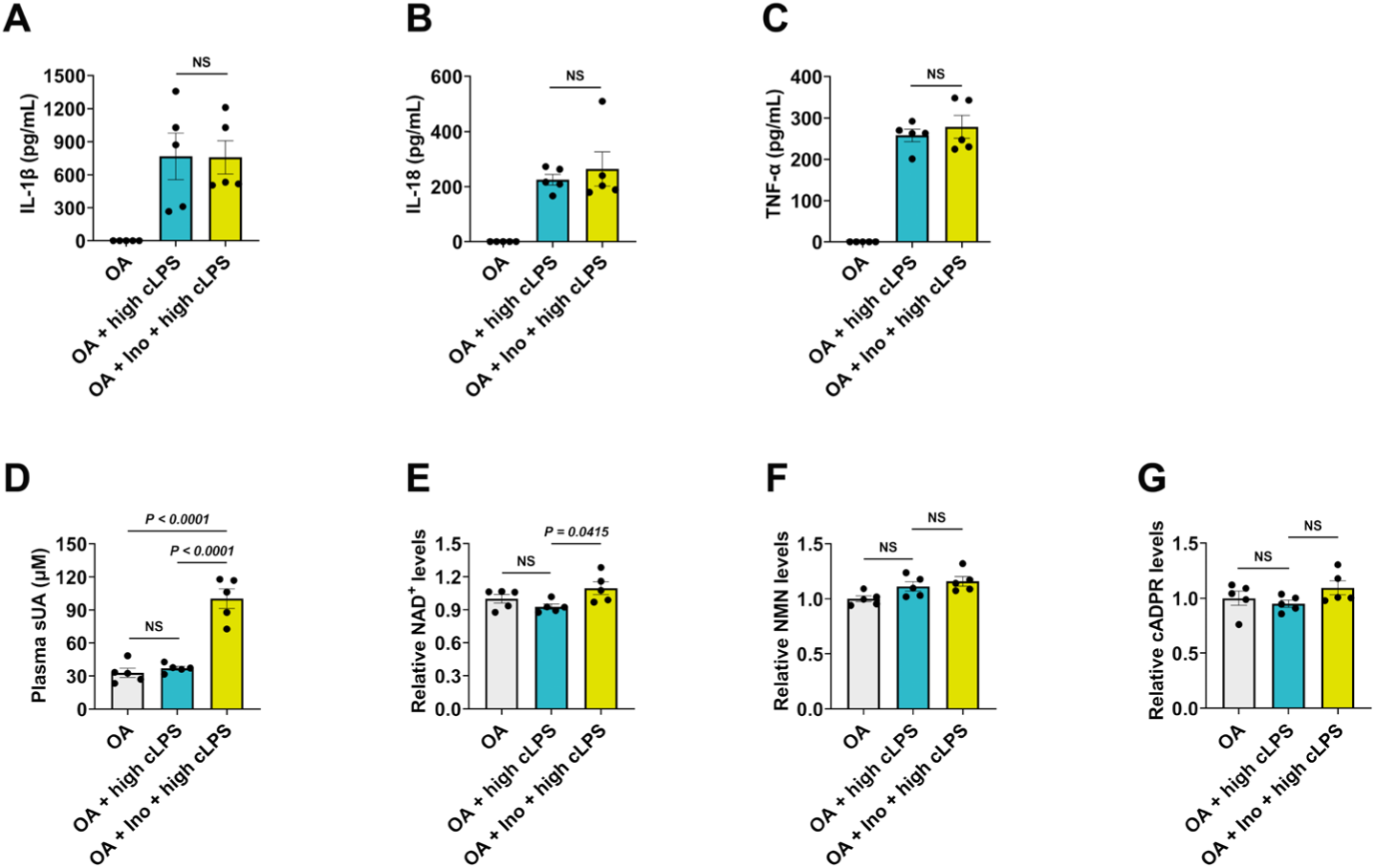
Moderate sUA supplementation fails to prevent high-dose cLPS-induced systemic inflammation. WT mice (10- to 12-week-old) received 1-day treatment of OA or OA plus inosine (Ino), 2 h after the last treatment, the mice were intraperitoneally stimulated with sterile PBS or cLPS (20 mg/kg) for 4 h (n = 5 mice per group). **(A-C)** Serum IL-1β (**A**), IL-18 (**B**), and TNF-α (**C**) were measured. **(D-G)** Plasma sUA (**D**), whole blood NAD^+^ (**E**), NMN (**F**), and cADPR (**G**) levels were measured. Data are mean ± s.e.m. Significance was tested using 1-way ANOVA with Tukey’s multiple comparisons test (**A**, and **C-G**) or Kruskal-Wallis test (**B**).

**Figure 4-figure supplement 5.**
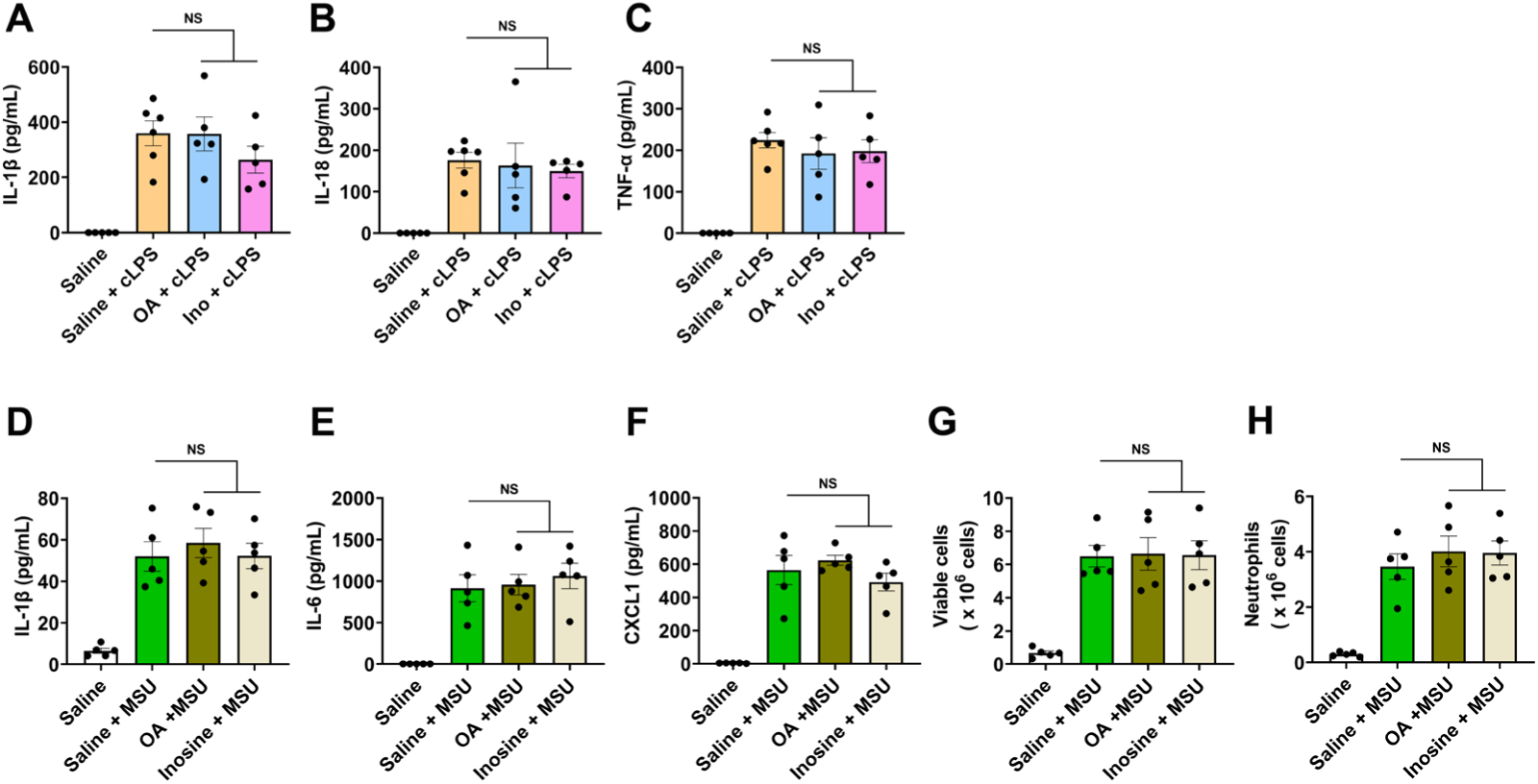
OA or inosine alone does not limit cLPS-induced systemic inflammation and MSU crystal-induced peritonitis. WT mice (10- to 12-week-old) received 1-day oral administration of saline, OA, or inosine (Ino), 2 h after the last treatment, the mice were intraperitoneally stimulated with sterile PBS, cLPS (2 mg/kg), or MSU crystals (2 mg/mouse) for 6 h. (**A-C)** Serum IL-1β (**A**), IL-18 (**B**), and TNF-α (**C**) levels in mice with cLPS-induced systemic inflammation were measured (n = 6 mice in Saline + cLPS group, n = 5 mice in other groups). (**D-H)** IL-1β (**D**), IL-6 (**E**), CXCL1 (**F**), and the number of viable cells (red blood cells excluded) (**G**) and neutrophils (**H**) in peritoneal lavage fluid from the mice with MSU crystal-induced peritonitis were measured (n = 5 mice per group). Data are mean ± s.e.m. Significance was tested using 1-way ANOVA with Tukey’s (**A, C-H**) multiple comparisons test, or Kruskal-Wallis test (**B**).

**Figure 4-figure supplement 6.**
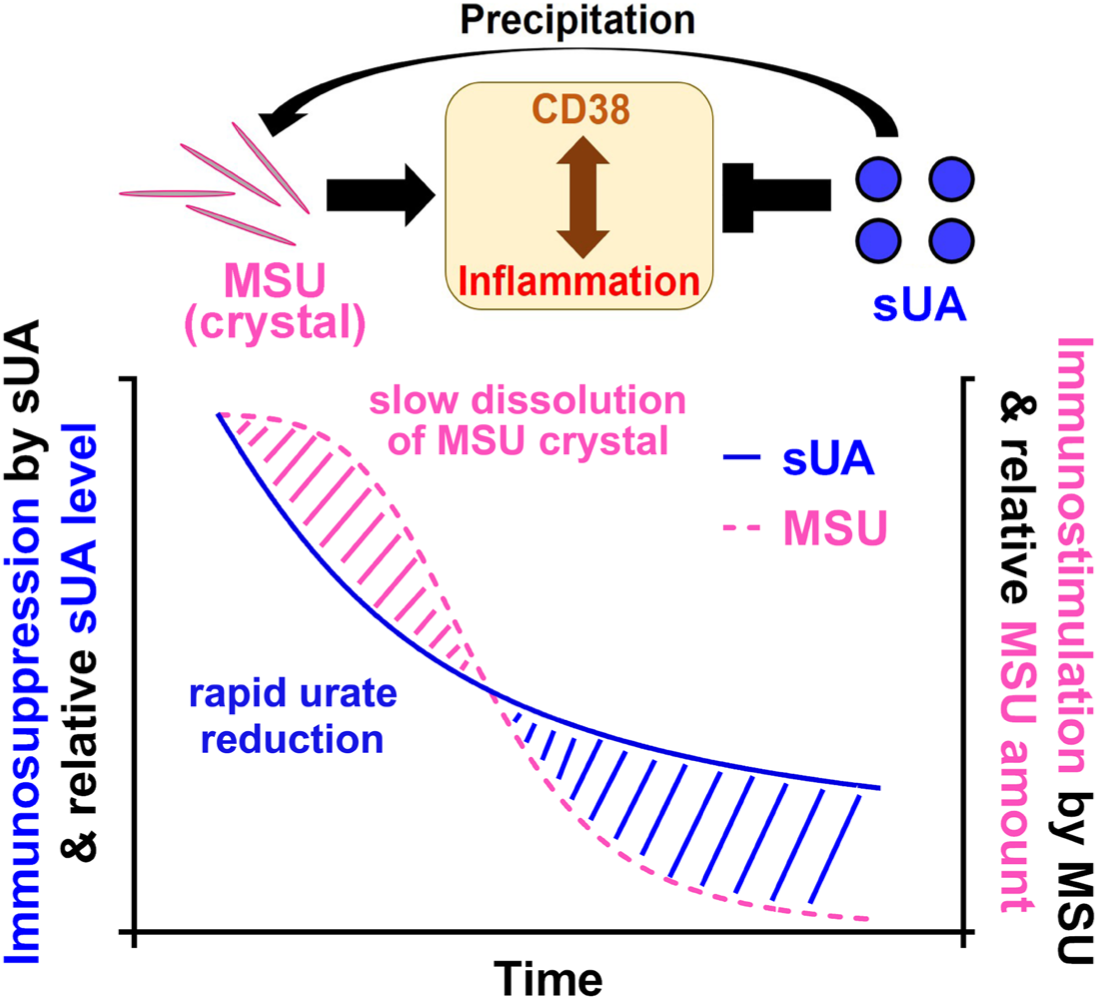
Potential mechanism of the paradox in gout therapy (urate-lowering therapy).

## References

Alabarse, P. G., Oliveira, P., Qin, H., Yan, T., Migaud, M., Terkeltaub, R., & Liu-Bryan, R. (2024). The NADase CD38 is a central regulator in gouty inflammation and a novel druggable therapeutic target. Inflamm Res, 73(5), 739–751. doi:10.1007/s00011-024-01863-y

Alberts, B. M., Barber, J. S., Sacre, S. M., Davies, K. A., Ghezzi, P., & Mullen, L. M. (2019). Precipitation of Soluble Uric Acid Is Necessary for In Vitro Activation of the NLRP3 Inflammasome in Primary Human Monocytes. J Rheumatol, 46(9), 1141–1150. doi:10.3899/jrheum.180855

Álvarez-Lario, B., & Macarrón-Vicente, J. (2010). Uric acid and evolution. Rheumatology (Oxford), 49(11), 2010–2015. doi:10.1093/rheumatology/keq204

Ames, B. N., Cathcart, R., Schwiers, E., & Hochstein, P. (1981). Uric acid provides an antioxidant defense in humans against oxidant- and radical-caused aging and cancer: a hypothesis. Proc Natl Acad Sci U S A, 78(11), 6858–6862. doi:10.1073/pnas.78.11.6858

Becker, M. A., Schumacher, H. R., Jr., Wortmann, R. L., MacDonald, P. A., Eustace, D., Palo, W. A.,… Joseph-Ridge, N. (2005). Febuxostat compared with allopurinol in patients with hyperuricemia and gout. N Engl J Med, 353(23), 2450–2461. doi:10.1056/NEJMoa050373

Blacher, E., Dadali, T., Bespalko, A., Haupenthal, V. J., Grimm, M. O., Hartmann, T.,… Levy, A. (2015). Alzheimer’s disease pathology is attenuated in a CD38-deficient mouse model. Ann Neurol, 78(1), 88–103. doi:10.1002/ana.24425

Boulieu, R., Bory, C., Baltassat, P., & Gonnet, C. (1983). Hypoxanthine and xanthine levels determined by high-performance liquid chromatography in plasma, erythrocyte, and urine samples from healthy subjects: the problem of hypoxanthine level evolution as a function of time. Anal Biochem, 129(2), 398–404. doi:10.1016/0003-2697(83)90568-7

Camacho-Pereira, J., Tarragó, M. G., Chini, C. C. S., Nin, V., Escande, C., Warner, G. M.,… Chini, E. N. (2016). CD38 Dictates Age-Related NAD Decline and Mitochondrial Dysfunction through an SIRT3-Dependent Mechanism. Cell Metab, 23(6), 1127–1139. doi:10.1016/j.cmet.2016.05.006

Cantor, J. R., Abu-Remaileh, M., Kanarek, N., Freinkman, E., Gao, X., Louissaint, A., Jr.,… Sabatini, D. M. (2017). Physiologic Medium Rewires Cellular Metabolism and Reveals Uric Acid as an Endogenous Inhibitor of UMP Synthase. Cell, 169(2), 258–272.e217. doi:10.1016/j.cell.2017.03.023

Chini, C. C. S., Peclat, T. R., Warner, G. M., Kashyap, S., Espindola-Netto, J. M., de Oliveira, G. C.,… Chini, E. N. (2020). CD38 ecto-enzyme in immune cells is induced during aging and regulates NAD(+) and NMN levels. Nat Metab, 2(11), 1284–1304. doi:10.1038/s42255-020-00298-z

Chini, E. N., Chini, C. C. S., Espindola Netto, J. M., de Oliveira, G. C., & van Schooten, W. (2018). The Pharmacology of CD38/NADase: An Emerging Target in Cancer and Diseases of Aging. Trends Pharmacol Sci, 39(4), 424–436. doi:10.1016/j.tips.2018.02.001

Chini, E. N., de Toledo, F. G., Thompson, M. A., & Dousa, T. P. (1997). Effect of estrogen upon cyclic ADP ribose metabolism: beta-estradiol stimulates ADP ribosyl cyclase in rat uterus. Proc Natl Acad Sci U S A, 94(11), 5872–5876. doi:10.1073/pnas.94.11.5872

Crawley, W. T., Jungels, C. G., Stenmark, K. R., & Fini, M. A. (2022). U-shaped association of uric acid to overall-cause mortality and its impact on clinical management of hyperuricemia. Redox Biol, 51, 102271. doi:10.1016/j.redox.2022.102271

Cutler, R. G., Camandola, S., Feldman, N. H., Yoon, J. S., Haran, J. B., Arguelles, S., & Mattson, M. P. (2019). Uric acid enhances longevity and endurance and protects the brain against ischemia. Neurobiol Aging, 75, 159–168. doi:10.1016/j.neurobiolaging.2018.10.031

Dalbeth, N., Gosling, A. L., Gaffo, A., & Abhishek, A. (2021). Gout. Lancet, 397(10287), 1843–1855. doi:10.1016/s0140-6736(21)00569-9

de Oliveira, G. C., Kanamori, K. S., Auxiliadora-Martins, M., Chini, C. C. S., & Chini, E. N. (2018). Measuring CD38 Hydrolase and Cyclase Activities: 1,N6-Ethenonicotinamide Adenine Dinucleotide (ε-NAD) and Nicotinamide Guanine Dinucleotide (NGD) Fluorescence-based Methods. Bio Protoc, 8(14), e2938. doi:10.21769/BioProtoc.2938

Deaglio, S., Aydin, S., Grand, M. M., Vaisitti, T., Bergui, L., D’Arena, G.,… Malavasi, F. (2010). CD38/CD31 interactions activate genetic pathways leading to proliferation and migration in chronic lymphocytic leukemia cells. Mol Med, 16(3-4), 87–91. doi:10.2119/molmed.2009.00146

Deaglio, S., Morra, M., Mallone, R., Ausiello, C. M., Prager, E., Garbarino, G.,… Malavasi, F. (1998). Human CD38 (ADP-ribosyl cyclase) is a counter-receptor of CD31, an Ig superfamily member. J Immunol, 160(1), 395–402. 10.4049/jimmunol.160.1.395

Dehlin, M., Jacobsson, L., & Roddy, E. (2020). Global epidemiology of gout: prevalence, incidence, treatment patterns and risk factors. Nat Rev Rheumatol, 16(7), 380–390. doi:10.1038/s41584-020-0441-1

Dogan, S., White, T. A., Deshpande, D. A., Murtaugh, M. P., Walseth, T. F., & Kannan, M. S. (2002). Estrogen increases CD38 gene expression and leads to differential regulation of adenosine diphosphate (ADP)-ribosyl cyclase and cyclic ADP-ribose hydrolase activities in rat myometrium. Biol Reprod, 66(3), 596–602. doi:10.1095/biolreprod66.3.596

Dudzinska, W., Lubkowska, A., Dolegowska, B., Safranow, K., & Jakubowska, K. (2010). Adenine, guanine and pyridine nucleotides in blood during physical exercise and restitution in healthy subjects. Eur J Appl Physiol, 110(6), 1155–1162. doi:10.1007/s00421-010-1611-7

Eells, J. T., & Spector, R. (1983). Purine and pyrimidine base and nucleoside concentrations in human cerebrospinal fluid and plasma. Neurochem Res, 8(11), 1451–1457. doi:10.1007/bf00965000

Escande, C., Nin, V., Price, N. L., Capellini, V., Gomes, A. P., Barbosa, M. T.,… Chini, E. N. (2013). Flavonoid apigenin is an inhibitor of the NAD+ ase CD38: implications for cellular NAD+ metabolism, protein acetylation, and treatment of metabolic syndrome. Diabetes, 62(4), 1084–1093. doi:10.2337/db12-1139

Fridovich, I. (1965). THE COMPETITIVE INHIBITION OF URICASE BY OXONATE AND BY RELATED DERIVATIVES OF S-TRIAZINES. J Biol Chem, 240, 2491–2494. 10.1016/S0021-9258(18)97351-5

Funaro, A., Horenstein, A. L., Calosso, L., Morra, M., Tarocco, R. P., Franco, L.,… Malavasi, F. (1996). Identification and characterization of an active soluble form of human CD38 in normal and pathological fluids. Int Immunol, 8(11), 1643–1650. doi:10.1093/intimm/8.11.1643

Glantzounis, G. K., Tsimoyiannis, E. C., Kappas, A. M., & Galaris, D. A. (2005). Uric acid and oxidative stress. Curr Pharm Des, 11(32), 4145–4151. doi:10.2174/138161205774913255

Gomez, G., & Sitkovsky, M. V. (2003). Differential requirement for A2a and A3 adenosine receptors for the protective effect of inosine in vivo. Blood, 102(13), 4472–4478. doi:10.1182/blood-2002-11-3624

Haberman, F., Tang, S. C., Arumugam, T. V., Hyun, D. H., Yu, Q. S., Cutler, R. G.,… Mattson, M. P. (2007). Soluble neuroprotective antioxidant uric acid analogs ameliorate ischemic brain injury in mice. Neuromolecular Med, 9(4), 315–323. doi:10.1007/s12017-007-8010-1

Hara-Yokoyama, M., Kukimoto, I., Nishina, H., Kontani, K., Hirabayashi, Y., Irie, F.,… Katada, T. (1996). Inhibition of NAD+ glycohydrolase and ADP-ribosyl cyclase activities of leukocyte cell surface antigen CD38 by gangliosides. J Biol Chem, 271(22), 12951–12955. doi:10.1074/jbc.271.22.12951

He, M., Chiang, H. H., Luo, H., Zheng, Z., Qiao, Q., Wang, L.,… Chen, D. (2020). An Acetylation Switch of the NLRP3 Inflammasome Regulates Aging-Associated Chronic Inflammation and Insulin Resistance. Cell Metab, 31(3), 580–591.e585. doi:10.1016/j.cmet.2020.01.009

Higashida, H., Egorova, A., Higashida, C., Zhong, Z. G., Yokoyama, S., Noda, M., & Zhang, J. S. (1999). Sympathetic potentiation of cyclic ADP-ribose formation in rat cardiac myocytes. J Biol Chem, 274(47), 33348–33354. doi:10.1074/jbc.274.47.33348

Hogan, K. A., Chini, C. C. S., & Chini, E. N. (2019). The Multi-faceted Ecto-enzyme CD38: Roles in Immunomodulation, Cancer, Aging, and Metabolic Diseases. Front Immunol, 10, 1187. doi:10.3389/fimmu.2019.01187

Hooftman, A., Angiari, S., Hester, S., Corcoran, S. E., Runtsch, M. C., Ling, C.,… O’Neill, L. A. J. (2020). The Immunomodulatory Metabolite Itaconate Modifies NLRP3 and Inhibits Inflammasome Activation. Cell Metab, 32(3), 468–478.e467. doi:10.1016/j.cmet.2020.07.016

Howles, S. A., & Thakker, R. V. (2020). Genetics of kidney stone disease. Nat Rev Urol, 17(7), 407–421. doi:10.1038/s41585-020-0332-x

Ichinose, W., Cherepanov, S. M., Shabalova, A. A., Yokoyama, S., Yuhi, T., Yamaguchi, H.,… Shuto, S. (2019). Development of a Highly Potent Analogue and a Long-Acting Analogue of Oxytocin for the Treatment of Social Impairment-Like Behaviors. J Med Chem, 62(7), 3297–3310. doi:10.1021/acs.jmedchem.8b01691

Ives, A., Nomura, J., Martinon, F., Roger, T., LeRoy, D., Miner, J. N., So, A. (2015). Xanthine oxidoreductase regulates macrophage IL1β secretion upon NLRP3 inflammasome activation. Nat Commun, 6, 6555. doi:10.1038/ncomms7555

Iwama, M., Kondo, Y., Shimokado, K., Maruyama, N., & Ishigami, A. (2012). Uric acid levels in tissues and plasma of mice during aging. Biol Pharm Bull, 35(8), 1367–1370. doi:10.1248/bpb.b12-00198

Jin, D., Liu, H. X., Hirai, H., Torashima, T., Nagai, T., Lopatina, O., Higashida, H. (2007). CD38 is critical for social behaviour by regulating oxytocin secretion. Nature, 446(7131), 41–45. doi:10.1038/nature05526

Joosten, L. A. B., Crişan, T. O., Bjornstad, P., & Johnson, R. J. (2020). Asymptomatic hyperuricaemia: a silent activator of the innate immune system. Nat Rev Rheumatol, 16(2), 75–86. doi:10.1038/s41584-019-0334-3

Kono, H., Chen, C. J., Ontiveros, F., & Rock, K. L. (2010). Uric acid promotes an acute inflammatory response to sterile cell death in mice. J Clin Invest, 120(6), 1939–1949. doi:10.1172/jci40124

Kutzing, M. K., & Firestein, B. L. (2008). Altered uric acid levels and disease states. J Pharmacol Exp Ther, 324(1), 1–7. doi:10.1124/jpet.107.129031

Kuwabara, M., Niwa, K., Ohtahara, A., Hamada, T., Miyazaki, S., Mizuta, E.,… Hisatome, I. (2017). Prevalence and complications of hypouricemia in a general population: A large-scale cross-sectional study in Japan. PLoS One, 12(4), e0176055. doi:10.1371/journal.pone.0176055

Kuzuya, M., Ando, F., Iguchi, A., & Shimokata, H. (2002). Effect of aging on serum uric acid levels: longitudinal changes in a large Japanese population group. J Gerontol A Biol Sci Med Sci, 57(10), M660–664. doi:10.1093/gerona/57.10.m660

Lai, J. H., Luo, S. F., Hung, L. F., Huang, C. Y., Lien, S. B., Lin, L. C.,… Ho, L. J. (2017). Physiological concentrations of soluble uric acid are chondroprotective and anti-inflammatory. Sci Rep, 7(1), 2359. doi:10.1038/s41598-017-02640-0

Lee, H. C. (2001). Physiological functions of cyclic ADP-ribose and NAADP as calcium messengers. Annu Rev Pharmacol Toxicol, 41, 317–345. doi:10.1146/annurev.pharmtox.41.1.317

Lee, H. C., Walseth, T. F., Bratt, G. T., Hayes, R. N., & Clapper, D. L. (1989). Structural determination of a cyclic metabolite of NAD+ with intracellular Ca2+-mobilizing activity. J Biol Chem, 264(3), 1608–1615. 10.1016/S0021-9258(18)94230-4

Lee, H. C., & Zhao, Y. J. (2019). Resolving the topological enigma in Ca(2+) signaling by cyclic ADP-ribose and NAADP. J Biol Chem, 294(52), 19831–19843. doi:10.1074/jbc.REV119.009635

Linnerz, T., Sung, Y. J., Rolland, L., Astin, J. W., Dalbeth, N., & Hall, C. J. (2022). Uricase-Deficient Larval Zebrafish with Elevated Urate Levels Demonstrate Suppressed Acute Inflammatory Response to Monosodium Urate Crystals and Prolonged Crystal Persistence. Genes (Basel), 13(12). doi:10.3390/genes13122179

Logan, J. A., Morrison, E., & McGill, P. E. (1997). Serum uric acid in acute gout. Ann Rheum Dis, 56(11), 696–697. doi:10.1136/ard.56.11.696a

Lomenick, B., Hao, R., Jonai, N., Chin, R. M., Aghajan, M., Warburton, S.,… Huang, J. (2009). Target identification using drug affinity responsive target stability (DARTS). Proc Natl Acad Sci U S A, 106(51), 21984–21989. doi:10.1073/pnas.0910040106

Lowell, C. A. (2022). Hyperuricemia reduces neutrophil function. Blood, 139(23), 3354–3356. doi:10.1182/blood.2022016275

Lu, N., Dubreuil, M., Zhang, Y., Neogi, T., Rai, S. K., Ascherio, A.,… Choi, H. K. (2016). Gout and the risk of Alzheimer’s disease: a population-based, BMI-matched cohort study. Ann Rheum Dis, 75(3), 547–551. doi:10.1136/annrheumdis-2014-206917

Luongo, T. S., Eller, J. M., Lu, M. J., Niere, M., Raith, F., Perry, C.,… Baur, J. A. (2020). SLC25A51 is a mammalian mitochondrial NAD(+) transporter. Nature, 588(7836), 174–179. doi:10.1038/s41586-020-2741-7

Ma, Q., Honarpisheh, M., Li, C., Sellmayr, M., Lindenmeyer, M., Böhland, C.,… Steiger, S. (2020). Soluble Uric Acid Is an Intrinsic Negative Regulator of Monocyte Activation in Monosodium Urate Crystal-Induced Tissue Inflammation. J Immunol, 205(3), 789–800. doi:10.4049/jimmunol.2000319

Ma, Q., Immler, R., Pruenster, M., Sellmayr, M., Li, C., von Brunn, A.,… Steiger, S. (2022). Soluble uric acid inhibits β2 integrin-mediated neutrophil recruitment in innate immunity. Blood, 139(23), 3402–3417. doi:10.1182/blood.2021011234

Martinez Molina, D., Jafari, R., Ignatushchenko, M., Seki, T., Larsson, E. A., Dan, C.,… Nordlund, P. (2013). Monitoring drug target engagement in cells and tissues using the cellular thermal shift assay. Science, 341(6141), 84–87. doi:10.1126/science.1233606

Martinon, F., Pétrilli, V., Mayor, A., Tardivel, A., & Tschopp, J. (2006). Gout-associated uric acid crystals activate the NALP3 inflammasome. Nature, 440(7081), 237–241. doi:10.1038/nature04516

Meyer, T., Shimon, D., Youssef, S., Yankovitz, G., Tessler, A., Chernobylsky, T.,… Mayo, L. (2022). NAD(+) metabolism drives astrocyte proinflammatory reprogramming in central nervous system autoimmunity. Proc Natl Acad Sci U S A, 119(35), e2211310119. doi:10.1073/pnas.2211310119

Mills, K. F., Yoshida, S., Stein, L. R., Grozio, A., Kubota, S., Sasaki, Y.,… Imai, S. I. (2016). Long-Term Administration of Nicotinamide Mononucleotide Mitigates Age-Associated Physiological Decline in Mice. Cell Metab, 24(6), 795–806. doi:10.1016/j.cmet.2016.09.013

Minhas, P. S., Liu, L., Moon, P. K., Joshi, A. U., Dove, C., Mhatre, S.,… Andreasson, K. I. (2019). Macrophage de novo NAD(+) synthesis specifies immune function in aging and inflammation. Nat Immunol, 20(1), 50–63. doi:10.1038/s41590-018-0255-3

Misawa, T., Takahama, M., Kozaki, T., Lee, H., Zou, J., Saitoh, T., & Akira, S. (2013). Microtubule-driven spatial arrangement of mitochondria promotes activation of the NLRP3 inflammasome. Nat Immunol, 14(5), 454–460. doi:10.1038/ni.2550

Murakami, T., Ockinger, J., Yu, J., Byles, V., McColl, A., Hofer, A. M., & Horng, T. (2012). Critical role for calcium mobilization in activation of the NLRP3 inflammasome. Proc Natl Acad Sci U S A, 109(28), 11282–11287. doi:10.1073/pnas.1117765109

Nishida, Y. (1991). Inhibition of lipid peroxidation by methylated analogues of uric acid. J Pharm Pharmacol, 43(12), 885–887. doi:10.1111/j.2042-7158.1991.tb03204.x

Oda, M., Satta, Y., Takenaka, O., & Takahata, N. (2002). Loss of urate oxidase activity in hominoids and its evolutionary implications. Mol Biol Evol, 19(5), 640–653. doi:10.1093/oxfordjournals.molbev.a004123

Partida-Sánchez, S., Cockayne, D. A., Monard, S., Jacobson, E. L., Oppenheimer, N., Garvy, B.,… Lund, F. E. (2001). Cyclic ADP-ribose production by CD38 regulates intracellular calcium release, extracellular calcium influx and chemotaxis in neutrophils and is required for bacterial clearance in vivo. Nat Med, 7(11), 1209–1216. doi:10.1038/nm1101-1209

Peclat, T. R., Shi, B., Varga, J., & Chini, E. N. (2020). The NADase enzyme CD38: an emerging pharmacological target for systemic sclerosis, systemic lupus erythematosus and rheumatoid arthritis. Curr Opin Rheumatol, 32(6), 488–496. doi:10.1097/bor.0000000000000737

Piedra-Quintero, Z. L., Wilson, Z., Nava, P., & Guerau-de-Arellano, M. (2020). CD38: An Immunomodulatory Molecule in Inflammation and Autoimmunity. Front Immunol, 11, 597959. doi:10.3389/fimmu.2020.597959

Rajman, L., Chwalek, K., & Sinclair, D. A. (2018). Therapeutic Potential of NAD-Boosting Molecules: The In Vivo Evidence. Cell Metab, 27(3), 529–547. doi:10.1016/j.cmet.2018.02.011

Scott, G. S., Spitsin, S. V., Kean, R. B., Mikheeva, T., Koprowski, H., & Hooper, D. C. (2002). Therapeutic intervention in experimental allergic encephalomyelitis by administration of uric acid precursors. Proc Natl Acad Sci U S A, 99(25), 16303–16308. doi:10.1073/pnas.212645999

Shi, Y. (2010). Caught red-handed: uric acid is an agent of inflammation. J Clin Invest, 120(6), 1809–1811. doi:10.1172/jci43132

Shi, Y., Evans, J. E., & Rock, K. L. (2003). Molecular identification of a danger signal that alerts the immune system to dying cells. Nature, 425(6957), 516–521. doi:10.1038/nature01991

Tarragó, M. G., Chini, C. C. S., Kanamori, K. S., Warner, G. M., Caride, A., de Oliveira, G. C.,… Chini, E. N. (2018). A Potent and Specific CD38 Inhibitor Ameliorates Age-Related Metabolic Dysfunction by Reversing Tissue NAD(+) Decline. Cell Metab, 27(5), 1081–1095.e1010. doi:10.1016/j.cmet.2018.03.016

Traut, T. W. (1994). Physiological concentrations of purines and pyrimidines. Mol Cell Biochem, 140(1), 1–22. doi:10.1007/bf00928361

Urano, W., Yamanaka, H., Tsutani, H., Nakajima, H., Matsuda, Y., Taniguchi, A.,… Kamatani, N. (2002). The inflammatory process in the mechanism of decreased serum uric acid concentrations during acute gouty arthritis. J Rheumatol, 29(9), 1950–1953. https://www.jrheum.org/content/29/9/1950

Wan, Q. L., Fu, X., Dai, W., Yang, J., Luo, Z., Meng, X.,… Zhou, Q. (2020). Uric acid induces stress resistance and extends the life span through activating the stress response factor DAF-16/FOXO and SKN-1/NRF2. Aging (Albany NY), 12(3), 2840–2856. doi:10.18632/aging.102781

Watanabe, S., Kang, D. H., Feng, L., Nakagawa, T., Kanellis, J., Lan, H.,… Johnson, R. J. (2002). Uric acid, hominoid evolution, and the pathogenesis of salt-sensitivity. Hypertension, 40(3), 355–360. doi:10.1161/01.hyp.0000028589.66335.aa

Wen, S., Arakawa, H., & Tamai, I. (2021). CD38 activation by monosodium urate crystals contributes to inflammatory responses in human and murine macrophages. Biochem Biophys Res Commun, 581, 6–11. doi:10.1016/j.bbrc.2021.10.010

Wen, S., Arakawa, H., & Tamai, I. (2024). Uric acid in health and disease: From physiological functions to pathogenic mechanisms. Pharmacol Ther, 256, 108615. doi:10.1016/j.pharmthera.2024.108615

Wu, X. W., Muzny, D. M., Lee, C. C., & Caskey, C. T. (1992). Two independent mutational events in the loss of urate oxidase during hominoid evolution. J Mol Evol, 34(1), 78–84. doi:10.1007/bf00163854

Yan, T. C. (2021). CD38 Inhibition Attenuates Monosodium Urate Crystal-induced Inflammation in Macrophages. University of California, San Diego. https://escholarship.org/uc/item/5nd3n13s

Zeidler, J. D., Kashyap, S., Hogan, K. A., & Chini, E. N. (2022). Implications of the NADase CD38 in COVID pathophysiology. Physiol Rev, 102(1), 339–341. doi:10.1152/physrev.00007.2021

Zhao, Y. J., Lam, C. M., & Lee, H. C. (2012). The membrane-bound enzyme CD38 exists in two opposing orientations. Sci Signal, 5(241), ra67. doi:10.1126/scisignal.2002700

Zielinska, W., Barata, H., & Chini, E. N. (2004). Metabolism of cyclic ADP-ribose: Zinc is an endogenous modulator of the cyclase/NAD glycohydrolase ratio of a CD38-like enzyme from human seminal fluid. Life Sci, 74(14), 1781–1790. doi:10.1016/j.lfs.2003.08.033

