## Supplementary material for "Functional identification of soluble uric acid as an endogenous inhibitor of CD38": Key Resources Table

| **Key Resources Table** | | | | |
| --- | --- | --- | --- | --- |
| **Reagent type (species) or resource** | **Designation** | **Source or reference** | **Identifiers** | **Additional information** |
| chemical compound, drug | Uric acid | Sigma-Aldrich | Cat#U0881 |  |
| chemical compound, drug | β-Nicotinamide Adenine Dinucleotide (NAD+) | Nacalai | Cat#24338-86 |  |
| chemical compound, drug | β-Nicotinamide mononucleotide (NMN) | BLD Pharmatech | Cat#BD116593 |  |
| chemical compound, drug | cADP-Ribose (cADPR) | Santa Cruz | Cat#sc-201512 |  |
| chemical compound, drug | FYU-981 | Fuji Yakuhin Co., Ltd. | N/A |  |
| chemical compound, drug | 78c | Sigma-Aldrich | Cat#5.38763 |  |
| chemical compound, drug | N-cyclohexyl benzamide | Sigma-Aldrich | Cat#R531332 |  |
| chemical compound, drug | Inosine | Nacalai | Cat#07139-42 |  |
| chemical compound, drug | Hypoxanthine | Sigma-Aldrich | Cat#H-9636 |  |
| chemical compound, drug | Xanthine | Sigma-Aldrich | Cat#X0626 |  |
| chemical compound, drug | Allantoin | Sigma-Aldrich | Cat#A-7878 |  |
| chemical compound, drug | Adenosine | FUJIFILM Wako | Cat#010-10491 |  |
| chemical compound, drug | Guanosine | FUJIFILM Wako | Cat#079-01111 |  |
| chemical compound, drug | Uracil | FUJIFILM Wako | Cat#212-00062 |  |
| chemical compound, drug | 1,3-dihydroimidazol-2-one | BLD Pharmatech | Cat#BD00733674 |  |
| chemical compound, drug | Oxypurinol | Sigma-Aldrich | Cat#O-6881 |  |
| chemical compound, drug | Caffeine | FUJIFILM Wako | Cat#031-06792 |  |
| chemical compound, drug | 1-methyluric acid | Sigma-Aldrich | Cat#M6885 |  |
| chemical compound, drug | 1,3-dimethyluric acid | Santa Cruz | Cat#sc-206240 |  |
| chemical compound, drug | 1,7-dimethyluric acid | Cayman Chemical | Cat#19584 |  |
| chemical compound, drug | 1,3,7-trimethyluric acid | Cayman Chemical | Cat#16949 |  |
| chemical compound, drug | 8-oxoguanine | Cayman Chemical | Cat#89290 |  |
| chemical compound, drug | Oxonic acid potassium salt | Sigma-Aldrich | Cat#156124 |  |
| chemical compound, drug | LPS from Escherichia coli 0111:B4 (crude LPS) | Sigma-Aldrich | Cat#L4130 |  |
| chemical compound, drug | Ultrapure LPS from Escherichia coli 0111:B4 | InvivoGen | Cat#tlrl-3pelps |  |
| chemical compound, drug | Phorbol 12-Myristate 13-Acetate | FUJIFILM Wako | Cat#162-23591 |  |
| chemical compound, drug | Nigericin | Sigma-Aldrich | Cat#N-7143 |  |
| chemical compound, drug | ATP | Oriental Yeast | Cat#45142000 |  |
| chemical compound, drug | Zymosan A from Saccharomyces cerevisiae | Sigma-Aldrich | Cat#Z4250 |  |
| chemical compound, drug | nicotinamide 1, N^6^-ethenoadenine dinucleotide (ε-NAD+) | Sigma-Aldrich | Cat#N2630 |  |
| chemical compound, drug | nicotinamide guanine dinucleotide (NGD) | Sigma-Aldrich | Cat#N5131 |  |
| peptide, recombinant protein | Recombinant human CD38 protein | Sino Biological | Cat#10818-H08H |  |
| peptide, recombinant protein | Recombinant human M-CSF protein | Proteintech Group | Cat#HZ-1192 |  |
| commercial assay or kit | Human IL-1β ELISA Kit | R & D Systems | Cat#DY201 |  |
| commercial assay or kit | Mouse IL-1β ELISA Kit | R & D Systems | Cat#SMLB00C |  |
| commercial assay or kit | Mouse IL-6 ELISA Kit | R & D Systems | Cat#DY406-05 |  |
| commercial assay or kit | Mouse IL-18 ELISA Kit | R & D Systems | Cat#DY7625-05 |  |
| commercial assay or kit | Mouse TNF-α ELISA Kit | R & D Systems | Cat#DY410-05 |  |
| commercial assay or kit | Mouse CXCL1 ELISA Kit | R & D Systems | Cat#DY453-05 |  |
| commercial assay or kit | Wright-Giemsa Stain Kit | ScyTek Laboratories | Cat#WGK-1 |  |
| commercial assay or kit | CycLex NAMPT Colorimetric Assay Kit Ver.2 | MBL | Cat#CY-1251V2 |  |
| commercial assay or kit | HT Universal Colorimetric PARP Assay Kit | R & D Systems | Cat#4677-096-K |  |
| cell line | A549 | ATCC | CCL-185 |  |
| cell line | THP-1 | ATCC | TIB-202 |  |
| cell line | ICR bone marrow-derived macrophages | This paper | N/A |  |
| strain, strain background | ICR mice | Japan SLC, Inc. | N/A |  |
| strain, strain background | CD38 KO ICR mice | Dr. Haruhiro Higashida, Kanazawa University | N/A |  |
| software, algorithm | Prism 9 | GraphPad Software | https://www.graphpad.com |  |
| software, algorithm | Labsolutions software (version 5.97) | Shimadzu | https://www.shimadzu.com/ |  |
| other | RF-6000 Spectrofluorophotometer | Shimadzu | https://www.shimadzu.com/ |  |
| other | LC-30A system | Shimadzu | https://www.shimadzu.com/ |  |
| other | LCMS-8050 | Shimadzu | https://www.shimadzu.com/ |  |
